# Programming isotype specific plasma cell differentiation

**DOI:** 10.1101/2021.08.31.458458

**Authors:** Brett W. Higgins, Andrew G. Shuparski, Karen B. Miller, Amanda M. Robinson, Louise J. McHeyzer-Williams, Michael G. McHeyzer-Williams

**Affiliations:** Department of Immunology and Microbial Sciences, The Scripps Research Institute, La Jolla, California 92037, USA

**Keywords:** Plasma Cell, Isotype, Class Switch Recombination, Germinal Center, T helper cell, single cell RNA-seq, B Cell, Antibody

## Abstract

Antibodies are produced across multiple isotypes with distinct properties that coordinate initial antigen clearance and confer long-term antigen-specific immune protection. Here, we interrogate the molecular programs of isotype-specific murine plasma cells (PC) following helper T cell dependent immunization and within established steady-state immunity. Using integrated single cell strategies, we reveal conserved and divergent components of the rapid effector phase of antigen-specific IgM^+^ versus inflammation modulating programs dictated by IgG2a/b^+^ PC differentiation. During antibody affinity maturation, the germinal center (GC) cycle imparts separable programs for post-GC inhibitory IgG1^+^ and inflammatory IgG2a/b^+^ PC to direct long-term cellular function. In the steady-state, two subsets of IgM^+^ and separate IgG2b^+^ PC programs clearly segregate from splenic IgA^+^ PC programs that emphasize mucosal barrier protection. These diverse isotype-specific molecular pathways of PC differentiation control complementary modules of antigen clearance and immune protection that could be selectively targeted for immunotherapeutic applications and vaccine design.

## INTRODUCTION

In response to T dependent (TD) antigen, antigen-presenting B cells are activated by follicular helper T cells (T_FH_)(*1, 2*), and undergo class switch recombination (CSR) through cytokine driven signaling cascade events(*3–5*). As CD4^+^ T helper cells and innate lymphoid cells (ILCs) are categorized into subsets based on discrete immune function(*6, 7*), B cells can be similarly assorted by antibody class(*8, 9*). IgM^+^ antibody is secreted as a pentamer for avidity-based antigen neutralization, complement activation or labeling for phagocyte uptake. Each secreted antibody elicits separable immune responses through class specific F_c_ receptor binding such as type 1 inflammation induction by IgG2a/b^+^ subclasses and type 2 inflammation reduction by IgG1^+^ antibody(*10–12*). IgA^+^ antibody can be uniquely produced as a dimer wrapped by a secretory component for translocation across epithelial layers to provide type 3 mucosal barrier defense and enable commensal microbiota tolerance(*13–15*). The role of PCs extends beyond antibody secretion as IgM^+^ and IgA^+^ PCs have been linked with interleukin (IL)-10, IL-17, IL-35, and tumor necrosis factor (TNF)-α secretory activity(*16–20*). Thus, PC immune effector activity is divided by antibody isotype; however, there remains little resolution of the divergent transcriptional control of immune module function across these class-specific PC compartments.

PC terminal differentiation requires expression of a conserved transcriptional program, guided by the upregulation of master transcriptional regulators Blimp-1, Xbp1 and IRF4 with concurrent downregulation of B cell lineage driving factors such as Bcl6, Pax5, Pu.1 and IRF8(*21–29*). During antigen clearance, an initial wave of PC differentiation produces an effector cohort that survives only days secreting low affinity antibody for antigen control and Fc receptor driven immune regulation. This early immune response module is influenced by T_FH_ cell driven CSR which produces varied PC antibody classes. Subsequently, B cells enter a geminal center (GC) reaction under the cognate direction of T_FH_ cells to undergo affinity maturation and exit as post-GC ‘memory’ PCs capable of long-term survival for durative antigen-specific protection(*30–34*). During this ‘memory’ phase of the immune response, T_FH_ cell directed GC cycling likely imprints supplementary molecular programming necessary for post-GC PC function and survival. It remains unclear the extent to which CSR and GC cycling impact the expression of ancillary molecular programs that divide these PC cohorts functionally at the single cell level.

Targeted immunization with model antigen and the unimmunized “steady-state” murine models provide access to varied classes of PC. TD antigen NP-KLH (4-hydroxy-3-nitrophenylacetyl conjugated to Keyhole Limpet Hemocyanin) synchronizes intact immune systems *in vivo* allowing access to pre-GC and follicular B cells for single cell molecular analysis (*35, 36*). Using NP-KLH and TLR4 agonist MPL as an adjuvant, we have detailed an antigen-specific IgG2a/b^+^ and IgG1^+^ dominated response with the effector pre-GC phase consisting of an unmutated PC compartment at day five, and a post-GC phase by day fourteen where all PCs exhibit somatic mutation(*8, 36–38*). In contrast, an unimmunized “steady-state” murine system continually exposed to environmental and gut microbiota antigen presents an ongoing, polyclonal immune response composed of IgM^+^, IgA^+^, and IgG^+^ B cell classes. As a result, the steady-state system provides the capacity to compare isotype-specific PC subsets distributed across multiple lymphoid organs(*13, 39*). Combining the use of these two models enables the interrogation of discrete molecular programming expressed by each class and subclass of PC.

Here we utilize a quantitative gene targeted RNA sequencing strategy(*40, 41*) to interrogate the underlying transcriptional heterogeneity of class specific PCs at the single cell level. We demonstrate that antigen-specific effector IgM^+^ and inflammatory IgG^+^ PC subclasses segregate transcriptionally with divergent molecular programs acquired during the GC reaction. This divergence was recapitulated in the steady state by IgM^+^, IgG2b^+^ and IgA^+^ PCs which exhibited unique and defining molecular programs that extended further into two phenotypically and transcriptionally distinct subpopulations within the IgM^+^ compartment. These studies indicate divergent transcriptional programs are imprinted according to broad immune response modules in both effector and post-GC pathways of PC differentiation. We propose that differential isotype-specific plasma cell programming can be selectively targeted for immune therapeutic modification and future vaccine design.

## RESULTS

### Antigen-driven segregation of isotype-specific PC differentiation

Acute response PC compartments express antigen-binding BCR following helper T cell dependent immunization at both the extrafollicular effector PC phase (Day 5) and the post-GC PC differentiation phase(*9, 23, 38*) (Day14; **Fig 1A-B; Supplemental Fig 1A-F**). These flow cytometry-based isolation strategies capture the majority of PC activity *in vivo* as seen by antigen-binding and isotype-specific ELISPOT (**Fig 1C**; **Supp Fig 1G-H**). Expression of both CD138 and intracellular Blimp-1 protein attest further to the identity of antigen-specific IgM^+^ and class-switched PC at the effector and post-GC ‘memory’ PC phase in the draining lymph nodes (LN) and bone marrow (BM) (**Fig 1D**; **Supp Fig 1D-E**). Cell cycle indicator Ki67 varied across antigen-specific PC compartments with highest levels at day 5 in both IgM^+^ and class-switched compartments (**Fig 1D**). Bimodal distributions of Ki67 at day 14 post-GC represents time since GC exit and terminal differentiation both locally and within the BM. As expected, expression of the B cell isoform of CD45, B220, and the BCR co-receptor CD19 decreased differentially by location and time but additionally by class, highlighting the potential for programmatic divergence *in vivo* (**Fig 1D**).

**Figure 1:**
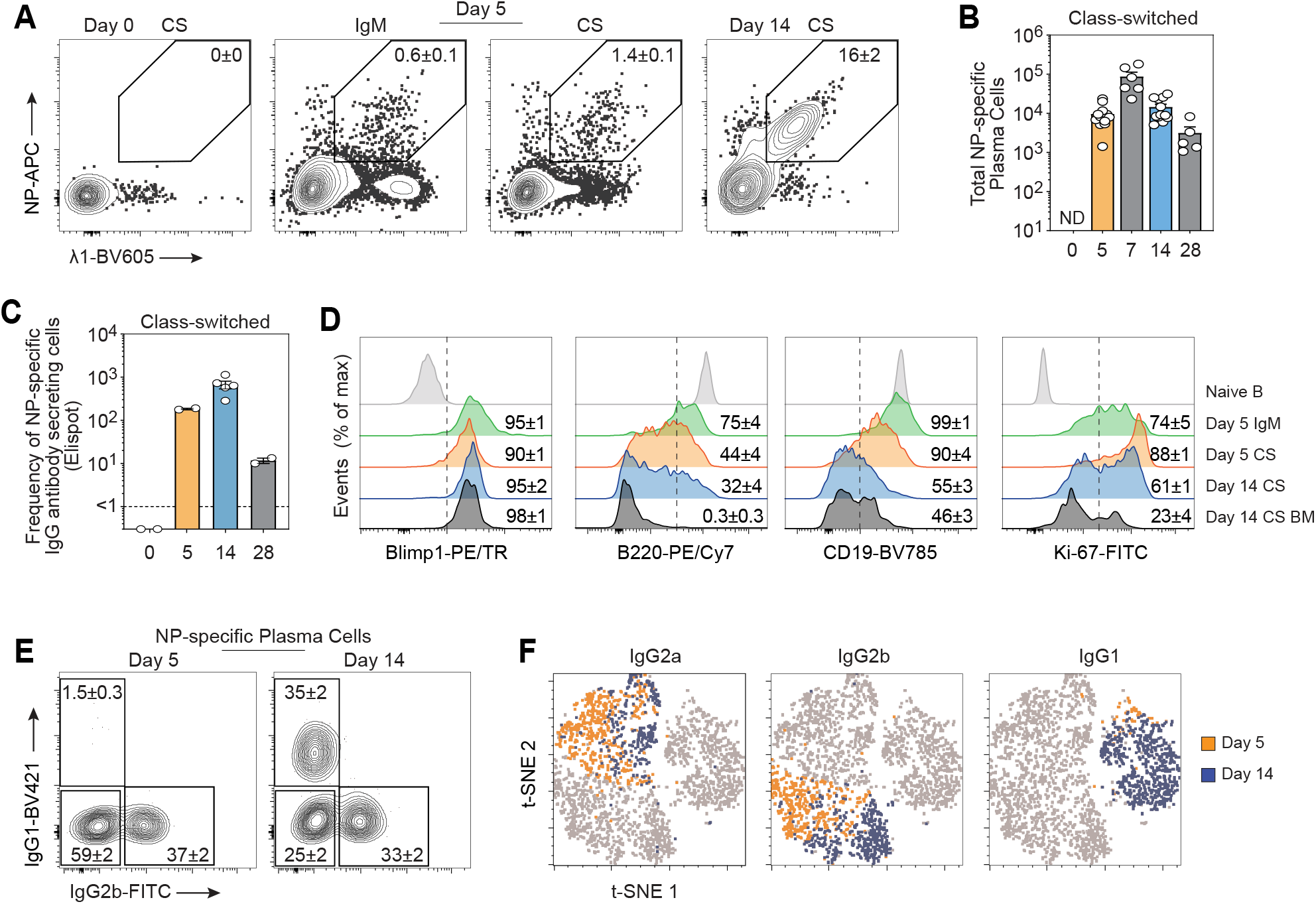
Antigen-driven segregation of isotype-specific PC differentiation. (**A**) Flow cytometry of class switched (CS, IgD^-^IgM^-^) or IgM^+^ (IgD-IgM^+^) antigen-specific (NP^+^λ1^+^) plasma cells (PC, Gr1^-^CD3e^-^IgD^-^CD138^+^) from draining lymph nodes (LN) before and 5 and 14 days after primary immunization with NP-KLH. Numbers in outlined areas indicate percent NP^+^λ1^+^ (mean± s.e.m.). (**B**) Quantification CS NP^+^λ1^+^ PCs from the dLN after primary immunization. ND, not detected. (**C**) Count per million frequency of CS NP^+^λ1^+^ PCs detected by Elispot at times (below graph) after immunization. (**D**) Intracellular expression of Blimp-1 and Ki-67 and surface expression of CD19 and B220 in NP^+^λ1^+^ PCs compared to naive B cells (CD19^+^IgD^+^IgM^+^λ1^+^CD138^-^NP^-^Gr1^-^CD3e^-^). Numbers indicate positive percentage expressing (mean ± s.e.m.). (**E**) Flow cytometry of IgG1^+^, IgG2b^+^ and IgG2a^+^ (IgG1^-^IgG2b^-^) NP^+^λ1^+^ PCs from d5 and d14. (**F**) Overlay of each PC class as shown in **E** (d5 in orange and d14 in blue) onto a tSNE (t-distributed stochastic neighbor embedding) plot generated using surface phenotype. (**A, B**; d0, d5, d14 n=12, d7 n=6, d28 n=5 mice per time point. **C**; 2-5 mice per time point; mean ± s.e.m. **D**; mean ± s.e.m., n=5 mice. **E**; mean ± s.e.m., d5 n=23, d14 n=18 mice. **F**; n=1534 cells at d5, n=1189 cells at d14 taken from 1 dLN at each time point, representative of n=12 mice)

IgM^+^ PCs represented only a minor fraction of responders to this TD antigen and MPL adjuvant. In contrast, among the IgG subclasses, type 1 inflammatory IgG2a^+^ and IgG2b^+^ emerged rapidly to dominate by day 5, accompanied by IgG1^+^ at day 7 with equivalent numbers of this type 2 anti-inflammatory isotype present post-GC by day 14 (**Fig 1E-F**; **Supp Fig 1F**). This antigen-specific PC differentiation model provides experimental access to early effector PCs expressing IgM^+^ and temporally and developmentally separate IgG^+^ subclasses to interrogate for transcriptional divergence at the single cell level.

### Divergent IgM and IgG effector PC Programs

As immunoglobulin mRNA makes up over 70% of sequenced cDNA species using global sequencing approaches in PCs(*29, 42, 43*), we developed a more targeted single cell RNA-seq protocol for these studies(*40, 41*). This quantitative and targeted RNA sequencing strategy (single cell qtSEQ) begins with high dimensional FACS sorted and indexed antigen-specific PC that are tracked and processed as single cells (**Fig 2A**). We currently target ∼500 gene products designed to exclude immunoglobulin genes but focus on expressed mRNA for transcriptional modifiers translocating to or residing in the nucleus (n=152), cell surface molecules (n=203) and cytoplasmic and secreted species (n=91 and n=51, respectively). Across this target set, barcoding was designed to identify reads from each individual sorted cell with the capacity to quantify per cell expression using unique molecular identifiers (UMIs) in a similar manner to global approaches (**Supplementary Fig 2A**). Dimensionality reduction of cell surface phenotype and gene expression distinguished the index sorted effector IgM^+^ and IgG^+^ PC compartments (**Fig 2B-C**). Relying on statistically significant divergence in gene expression, expression fold change (**Fig 2D-F**) and averaged heatmap representations of scaled signals in heatmaps (**Fig 2G**), we establish multiple isotype-segregating components of the IgM^+^ and class-switched antigen-specific effector PC transcriptional program.

**Figure 2:**
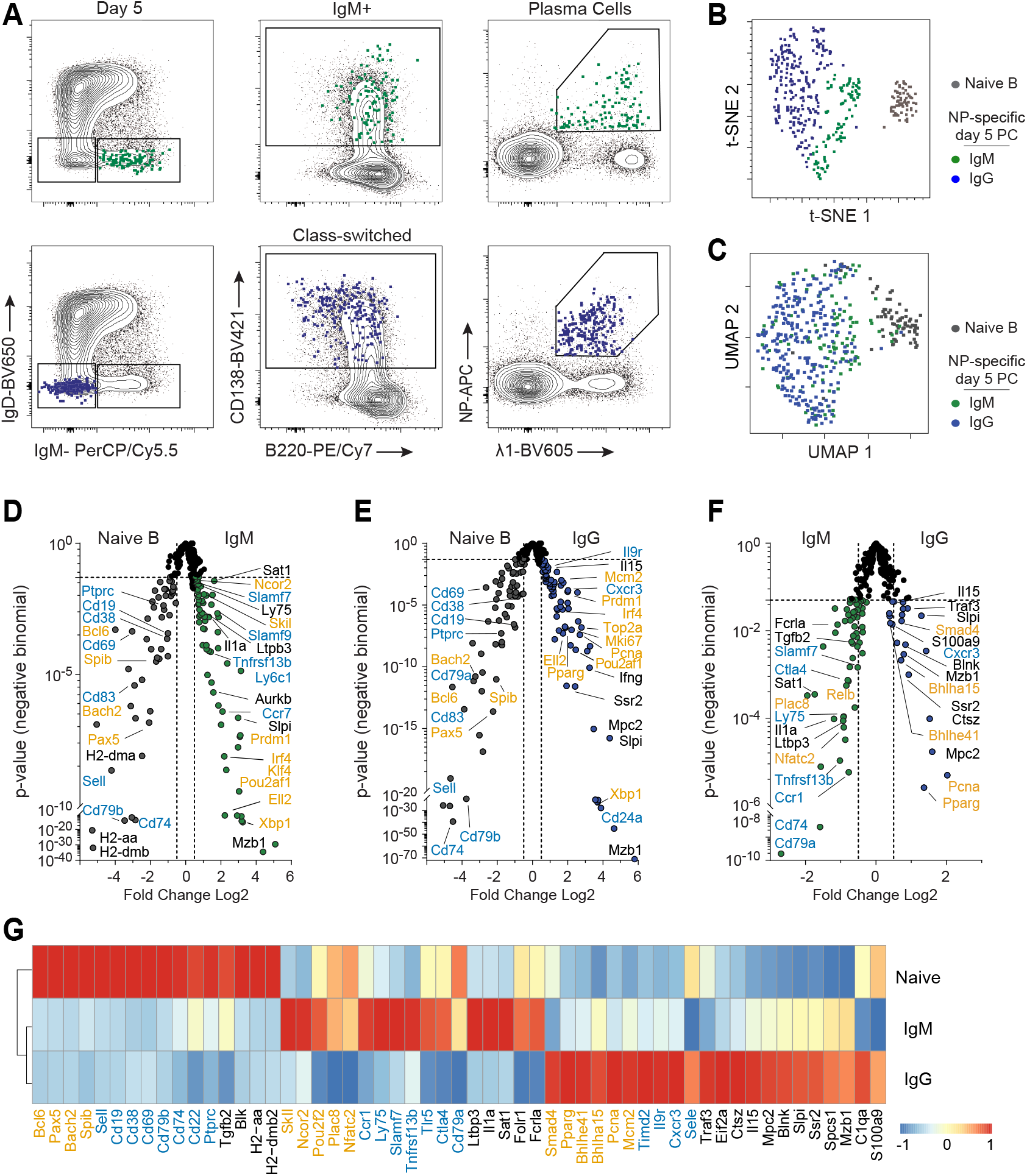
Divergent IgM and IgG effector PC Programs. (**A**) Flow cytometry of index sorted antigen specific IgM^+^ (top, in green) and IgG^+^ (bottom, in blue) PCs overlaid on contour plots of total populations. (B) Dimensional reduction on index sorted PCs based on surface phenotype using tSNE (t-distributed stochastic neighbor embedding) or (**C**) based on gene expression distribution using uniform manifold approximation and projection (UMAP). (**D**) Differences in gene expression plotted by statistical significance and presented as volcano plots comparing naive B cells (CD19^+^IgD^+^IgM^+^λ1^+^CD138^-^NP^-^Gr1^-^CD3e^-^) to antigen specific IgM^+^ PCs to (**E**) IgG^+^ PCs, or between (**F**) IgM^+^ and IgG^+^ PCs. (**G**) Pseudobulk heatmap representation of differentially expressed genes from **D-F** using average gene expression values in each cell population. The color scale is based on z-score distribution. For **D-G**, index sorted cells n=109 IgM^+^, 269 IgG^+^, 73 naive B cells; genes labeled in orange represent transcriptional modifiers translocating to or residing in the nucleus, genes labeled in teal represent surface/membrane genes and genes in black represent cytosolic and secreted genes.

The loss of central naive B cell transcriptional programs (*Bcl6, Pax5*, *Bach2*, *Spib*) were a cardinal feature of early effector IgM^+^ PC differentiation (**Fig 2D; Supplementary Fig 2B**). These substantial changes in transcriptional regulation were accompanied by increased PC lineage drivers (e.g., *Prdm1*, *Irf4*, *Xbp1*, *Pou2af1*) with some factors expressed significantly higher in extrafollicular IgM^+^ effector PC compared to their class-switched counterpart (**Fig 2D,E; Supplementary Fig 2C**). Multiple other transcriptional modifiers were differentially expressed at higher (*Skil, Ncor2, Pou2f2, Plac8, Nfatc2*) or lower levels (*Smad4, Traf3*, *Pparg, Bhlha15* and *Bhlhe41*) allow separation of the central programs of IgM^+^ from IgG^+^ class-switched PC at the effector stage (**Fig 2F,G**). IgM^+^ PCs may express ancillary effector functions related to differentially expressed secretory molecules (e.g., *Tgfb2, Ltbp3, Il1a*) in contrast to IgG^+^ PC from this stage (e.g., *Il15, Slpi*, *S100a9;* **Fig 2F,G**). Finally, there are significant differences in expression of surface effector molecules including chemokine receptors and cell adhesion mediators between PC subsets (e.g., *Ccr1*, *Tnfrsf13b, Slamf7 Ctla4, Cd79a* in IgM^+^*; Timd2, Il9r, Cxcr3, Sele* in IgG; **Fig 2G**). This programmatic heterogeneity can markedly impact class specific BCR responsiveness, immunomodulation, migratory behavior, and survival requirements. Together, IgM^+^ and IgG^+^ effector PC compartments demonstrate divergent transcriptional programs with evidence for separable secretory and surface modifiers that direct class-specific immune function.

### Inflammatory subclass IgG effector PC programs

Closer scrutiny of IgG2a^+^ and IgG2b^+^ effector PC (**Fig 3A; Supplementary Fig 3**) revealed further significant transcriptional differences between the cells expressing these early inflammatory mediating antibody subclasses (**Fig 3B-D**). Expression of major transcriptional regulators are significantly higher in IgG2a^+^ PC (*Foxj1, Klf6, Klf7, Bcl6b* and *Irf1*) in contrast to IgG2b^+^ PCs (*Pparg, Bhlha15, Bhlhe41, Xbp1, Ikzf1*; **Fig 3D,E**). Differential upregulation of secretory molecules that have been linked with eliciting inflammatory responses (*Il22, S100a9, Gzma* in IgG2a^+^; *Il15* in IgG2b^+^) as well as surface expressed proteins (*Tnfrsf1b,Tnfsf9, Tnfsf14, Itgb2* in IgG2a^+^; *Havcr2* in IgG2b^+^) further distinguished a separable IgG2a^+^ transcriptional module from IgG2b^+^ PCs. Consistent with the inflammatory functions of IgG2a/b subclass Cxcr3 is expressed highly compared to IgM and validated with protein expression. Surface expression of MHC-II by effector PCs suggested conserved retention of antigen presentation capacity, though differential Cxcr3 and B220 expression emphasized shared programmatic components of IgG^+^ subclasses that still differed from IgM^+^ PCs (**Fig 3F-H**). The separation of antigen-specific effector PCs into IgG2a^+^ and IgG2b^+^ subclasses emphasizes the existence of dichotomous subclass-linked molecular programming with evidence for heterogenous inflammatory response control.

**Figure 3:**
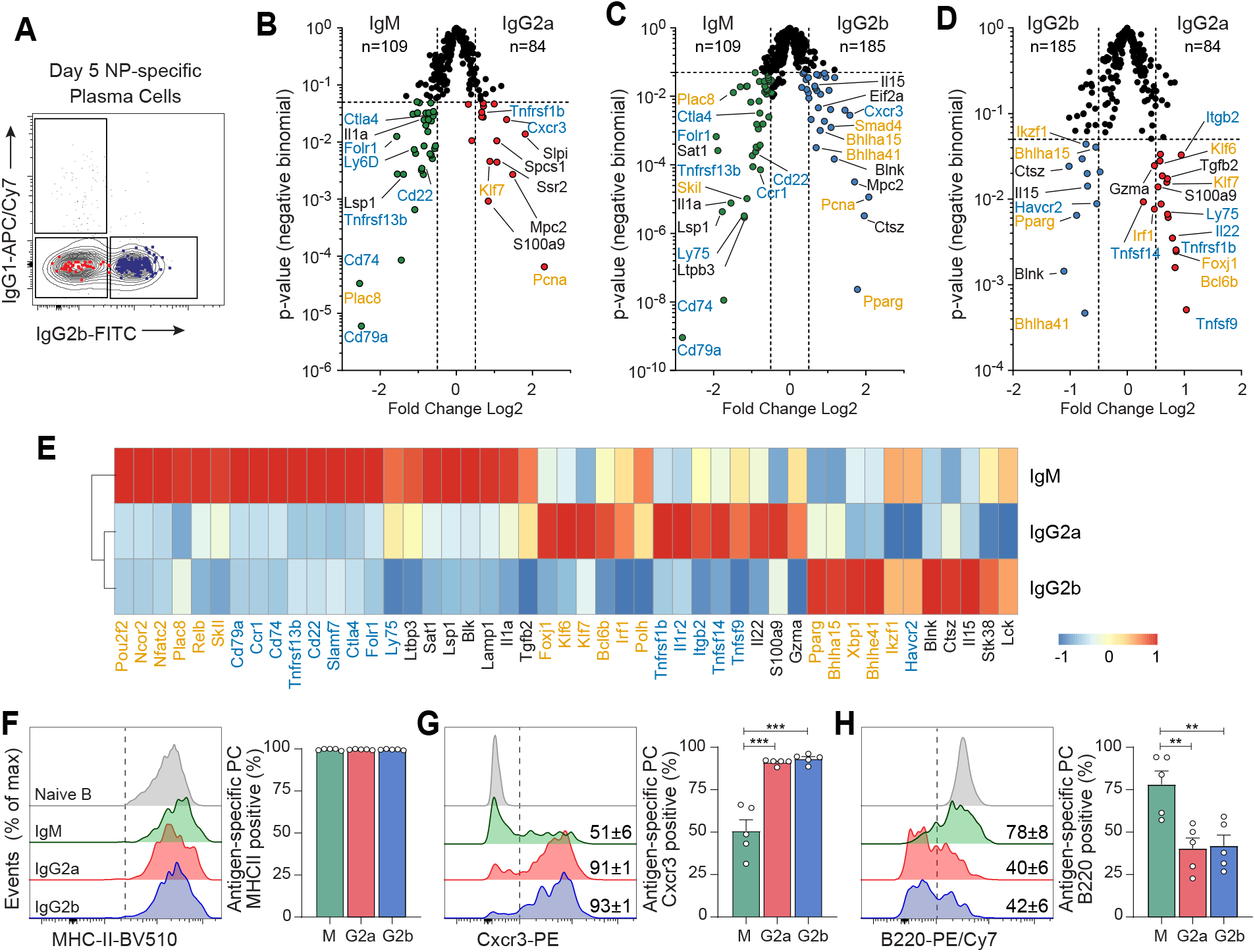
Inflammatory subclass IgG effector PC programs. (**A**) Flow cytometry of index sorted CS NP^+^λ1^+^ PCs (as in Fig 2A) split by surface IgG1^+^ and IgG2b^+^ expression. (**B-D**) Differences in gene expression plotted by statistical significance and presented as volcano plots comparing d5 antigen-specific IgM^+^, IgG2a^+^, and IgG2b^+^ PCs. (**E**) Pseudobulk heatmap representation of differentially expressed genes from **B-D**. The color scale is based on z-score distribution. (**F-H**) Surface protein expression of MHC-II, Cxcr3 and B220 from antigen-specific PCs split by class compared to naive B cells (CD19^+^IgD^+^IgM^+^λ1^-^CD138^-^NP^-^Gr1^-^CD3e^-^). Numbers indicate the percentage of cells expressing each surface marker (mean ± s.e.m., n=5; ** p <0.01, *** p<0.001, 2 tailed unpaired Students *t-*test). For index sorted cells, n=109 IgM^+^, 185 IgG2b^+^, 84 IgG2a^+^ PCs. For **B-E**, genes labeled in orange represent transcriptional modifiers translocating to or residing in the nucleus, genes labeled in teal represent surface/membrane genes and genes in black represent cytosolic and secreted genes.

### Divergent isotype-specific post-GC PC programs

The GC cycle is fully operative by day 14 of this immune response producing antigen-specific memory B cells and post-GC PCs that express affinity-matured BCR(*8, 35–38, 44–47*). This established GC reaction produces equivalent numbers of anti-inflammatory IgG1^+^ post-GC PC and their inflammatory IgG2a^+^ and IgG2b^+^ post-GC PC counterparts (**Fig 1E; Fig 4A-B)**. Compared to naïve B cells, post-GC IgG1^+^ PCs upregulated master transcriptional regulators involved in cell cycling, antibody secretion and UPR control (e.g., *Ube2c, Mki67, E2f2, Ell2, Xbp1, Atf4, Bhlha15*), cytokine linked secretory components (e.g., *Ifng, Il15, Il22*) and surface molecules (e.g., *Cd44, Itgal, Cd93, Cd28, Cd80, Cd276, Cd9*) that direct long-term cellular function (**Fig 4C-E**).

**Figure 4:**
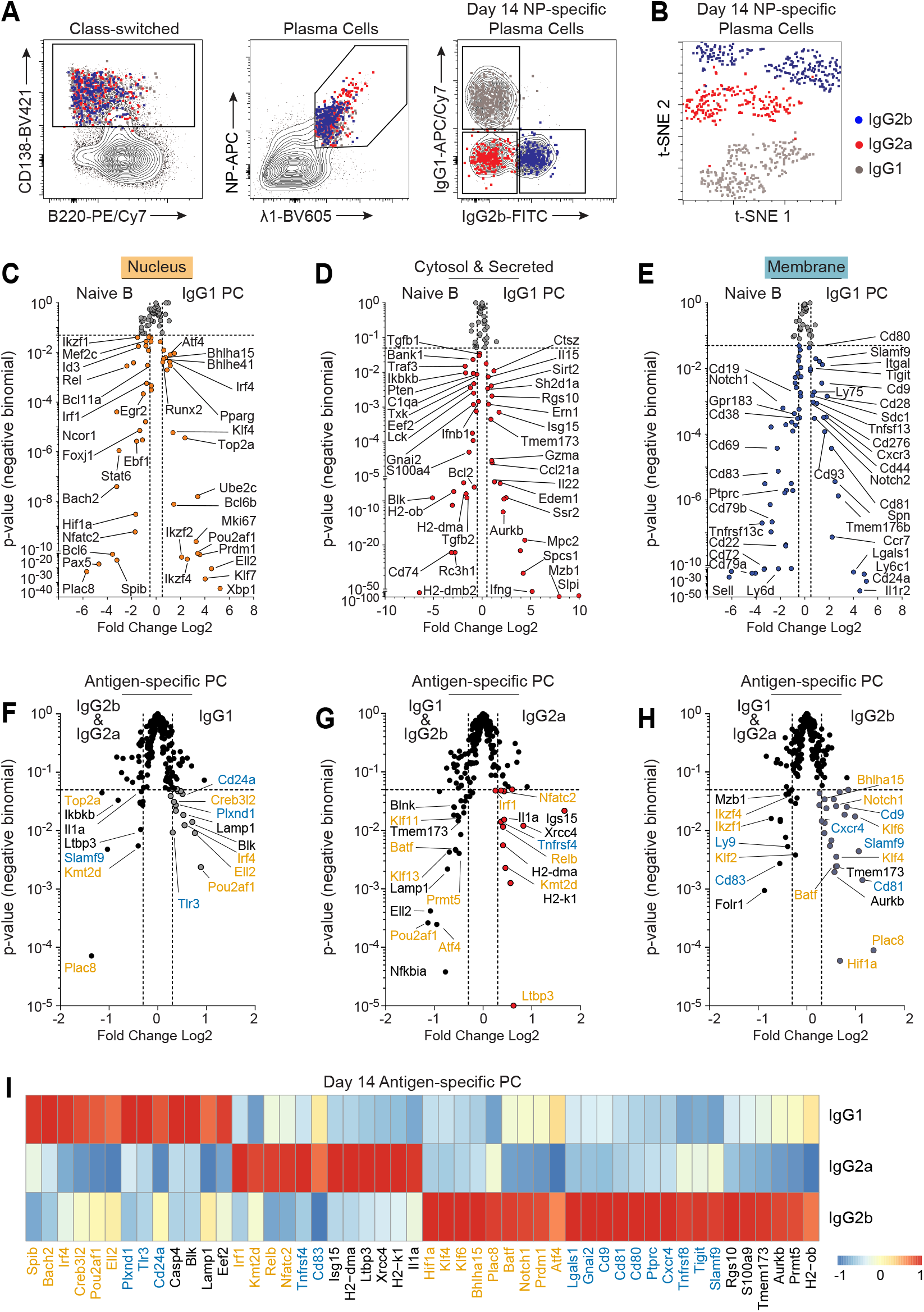
Divergent isotype-specific post-GC PC programs. (**A**) Index sorted antigen specific IgG1^+^, IgG2b^+^, and IgG2a^+^ PCs overlaid on contour plots of total populations. (**B**) Dimensional reduction based on surface phenotype of index sorted PCs using tSNE (t-distributed stochastic neighbor embedding) (n=290 IgG1^+^, n=292 IgG2b^+^, n=228 IgG2a^+^ PCs). (**C-E**) Differences in gene expression plotted by statistical significance and presented as volcano plots comparing naive B cells (CD19^+^IgD^+^IgM^+^Lambda1^+^CD138^-^NP^-^Gr1^-^CD3e^-^) and antigen specific IgG1^+^ PCs separated by gene categorization. (**F-H**) Differences in gene expression plotted by statistical significance and presented as volcano plots comparing antigen specific IgG1^+^ (**D**), IgG2b^+^ (**E**), and IgG2a^+^ (**F**) PCs compared to all other post-GC PCs. (**I**) Pseudobulk heatmap representation of differentially expressed genes from **F-H** using average gene expression values. The color scale is based on z-score distribution. (n=78 naïve, n=290 IgG1^+^, n=292 IgG2b^+^, n=228 IgG2a^+^ PCs). For **F-H**, genes labeled in orange represent transcriptional modifiers translocating to or residing in the nucleus, genes labeled in teal represent surface/membrane genes and genes in black represent cytosolic and secreted genes.

Comparison between post-GC PCs indicated IgG1^+^ post-GC PCs expressed significantly higher levels of central transcriptional regulators (*Spib, Bach2*, *Pou2af1, Creb3l2*, *Irf4, Ell2)* that would coordinate separable programs upon GC exit. In contrast, IgG2a^+^ post-GC PCs differentially expressed a separate set of regulators *(Irf1, Kmt2d, Relb, Nfatc2*) that segregate further against IgG2b^+^ post-GC PC (*Hif1a, Klf4, Klf6, Bhlha15, Plac8, Batf, Notch1, Prdm1* and *Atf4*) (**Fig 4F-H**). Surface expressed, cytosolic and secreted components of these programs begin to indicate the impact of the transcriptional differences on subclass specific post-GC PC function. IgG1^+^ post-GC PC differentially expressed multiple molecules with capacity to elicit divergent function (*Plxnd1, Tlr3, Cd24a, Blk, Lamp1;* **Fig 4F,I**); while IgG2a^+^ post-GC PCs exhibited unique gene upregulation including a set of secretory molecules (*Isg15, Ltbp3, Il1a*) associated with directing a range of inter-cellular functions (**Fig 4I,G**). In the context of the targeted gene set, IgG2b^+^ post-GC B cells differentially expressed multiple segregated programs, notably a series of cellular adhesion molecules (*Cd9, Cd81, Cd80, Ptprc, Tigit, Slamf9;* **Fig 4H,I**). These divergent transcriptional changes remain prominent and persistent even following the complex behavior of somatic hypermutation and antigen-driven selection that is paramount in the GC reaction.

As described above, we used Ki67 as a marker of recent cell cycling and hence as a proxy for recent GC cycle exit in post-GC PCs. We utilized index tracing to divide post-GC PCs by *Mki67* expression and determined that each IgG subclass contained equal frequencies of *Mki67*^+^ cells (**Supplementary Fig 4B**). As expected of cells more recently cycling, *Mki67*^+^ PCs were more transcriptionally active with enrichment of programs associated with apoptosis, cell division and replication compared to the *Mki67*^-^ population (**Supplementary Fig 4C,D**).

### Separable inflammatory post-GC PC programs

While post-GC PCs retained expression of MHC-II similar to effector PCs, they exhibit differential expression of CXCR3 and FAS (**Fig 5A**). Furthermore, gene program driven dimensionality reduction separates effector and post-GC PCs (**Fig 5B**). Through comparison of inflammatory class effector PCs to post-GC PCs, we can establish unique molecular programs imprinted by GC cycling (**Fig 5C-D**). Dividing our analysis into IgG2a^+^ and IgG2b^+^ subclasses resolved subclass specific transcriptional programs that emerge post-GC (**Fig 5E-G**). The reliance on highly influential modules of transcriptional regulation at day 5 (*Myc, Foxj1, Pcna, Notch2* in IgG2a*; E2f2, Ciita, Bhlha15, Pparg, Akt1, Smad4, Irf4, Ube2c* in IgG2b^+^) was significantly shifted by day 14 in the post-GC PC compartment, which upregulated a separate series of nuclear factors (*Relb, Kmt2d* in IgG2a^+^; *Klf7, Skil* in both; *Stat6, Mef2c* in IgG2b^+^) (**Fig 5G**).

**Figure 5:**
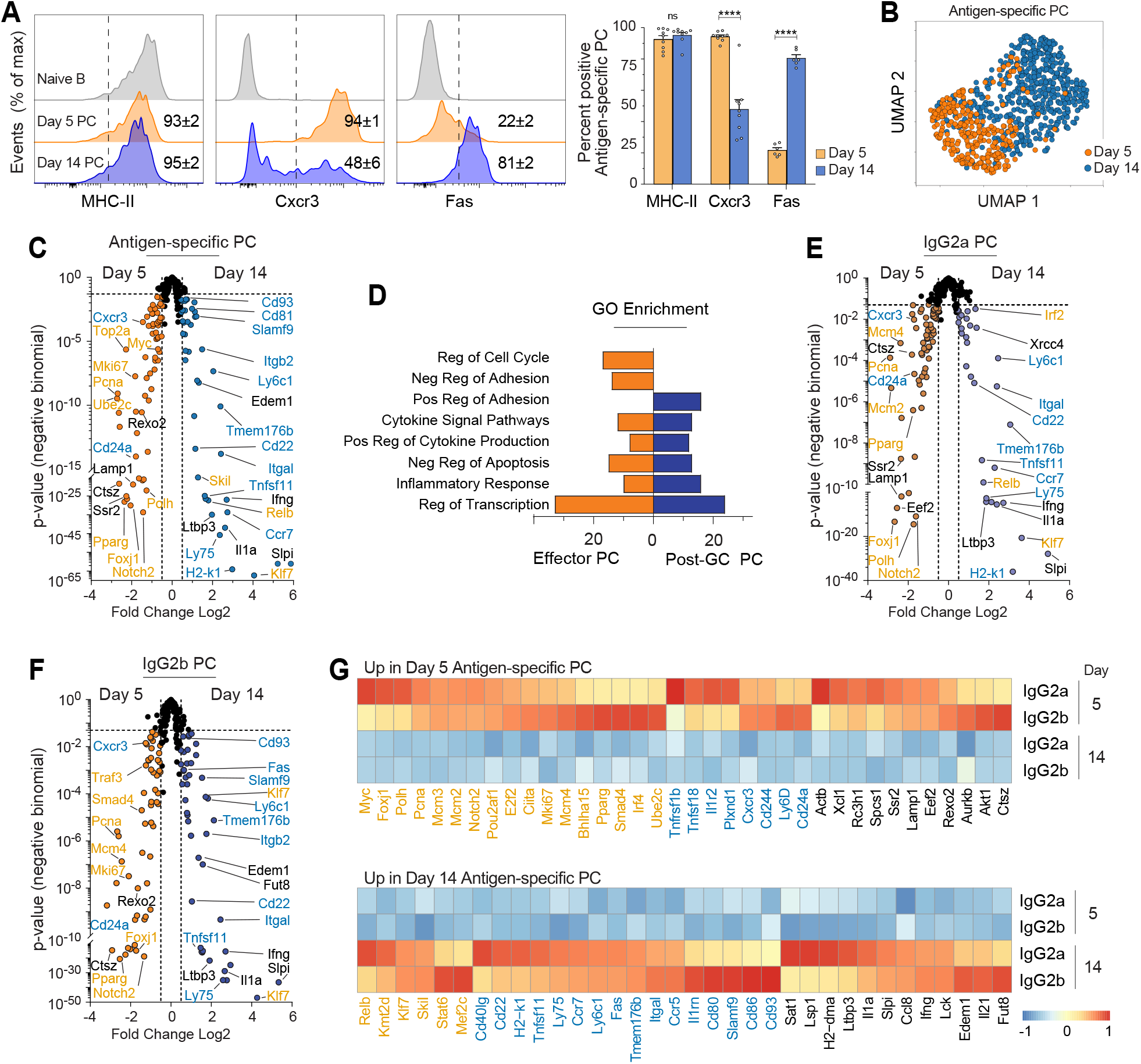
Separable inflammatory post-GC PC programs. (**A**) Surface protein expression of MHC-II, Cxcr3 and Fas in antigen specific d5 and d14 PCs compared to naive B cells. Numbers indicate percentage of cells expressing each marker. (mean ± s.e.m., n=6-9 mice, **** p<0.0001, ns, not significant; 2 tailed unpaired Students *t-*test). (**B**) UMAP reduction of antigen specific d5 and d14 PCs using gene expression (n=291 d5 PCs, n=520 d14 PCs). (**C**) Differences in gene expression plotted by statistical significance and presented as a volcano plot comparing d5 and d14 antigen specific PCs (IgG2a^+^ and IgG2b^+^ only). (**D**) The number of genes upregulated aligning to the indicated GO pathway showing d5 PCs in orange and d14 PCs in blue. (Cell cycle, GO:0051726; Neg Adhesion, GO:0007162; Pos Adhesion, GO:0022409; Cytokine Sig Pathway, GO 0019221; Pos Cytokine Production, GO:0001819; Neg Apoptosis, GO:0043066; Inflammatory resp, GO:0006954; Transcription, GO:0006355) (**E, F**) Differences in gene expression plotted by statistical significance and presented as a volcano plot comparing d5 and d14 antigen specific PCs split by IgG2b^+^ (**E**) or IgG2a^+^ (**F**). (**G**) Pseudobulk heatmap representation of differentially expressed genes between d5 and d14 PCs split by subclass. The color scale is based on z-score distribution. Genes labeled in orange represent transcriptional modifiers translocating to or residing in the nucleus, genes labeled in teal represent surface/membrane genes and genes in black represent cytosolic and secreted genes.

Beyond transcriptional regulation there were significant changes in expression of important cell membrane expressed regulators of immune function which extended further to segregation by antibody subclass; we revealed changes from day 5 effector PCs (e.g., *Tnfrsf1b, Tnfsf18, Il1r2,* in IgG2a; *Cxcr3, Cd244, Ly6D* and *Cd24a* in IgG2b) to post-GC PCs (e.g., *Cd40l, Cd22, Tnfsf11* in IgG2a; *Il1rn, Cd80, Slamf9, Cd86, Cd93* in IgG2b). Post-GC PCs also upregulated secretory factors (*Ltbp3*, *Il1a*, *Slpi, Ccl8*, *Ifng, Il21*) not represented in effector PC compartments. These subclass specific components of post-GC changes, including chemokine receptors, B7 family receptors, and adhesion molecules suggest shifts in cellular function imprinted by the GC program prior to post-GC PC differentiation.

### Divergent IgM and IgG2b PC programs in the steady state

Next, we chose to broaden our molecular analyses to the range of isotype-specific PC found in the steady-state spleen. While CD138 is a reliable PC marker in the context of antigen-binding responses, Blimp-1 reporter expression was used in these steady-state studies. High levels of Blimp-YFP in IgM^+^ or class-switched B cells served to identify antibody-secreting PC of separable isotype (**Fig 6A**; **Supplementary Fig 5A-B**). The IgM^+^ and IgG2b^+^ subsets broadly separated in UMAP clusters based on the full range of gene expression from qtSEQ analysis (**Fig 6B**).

**Figure 6:**
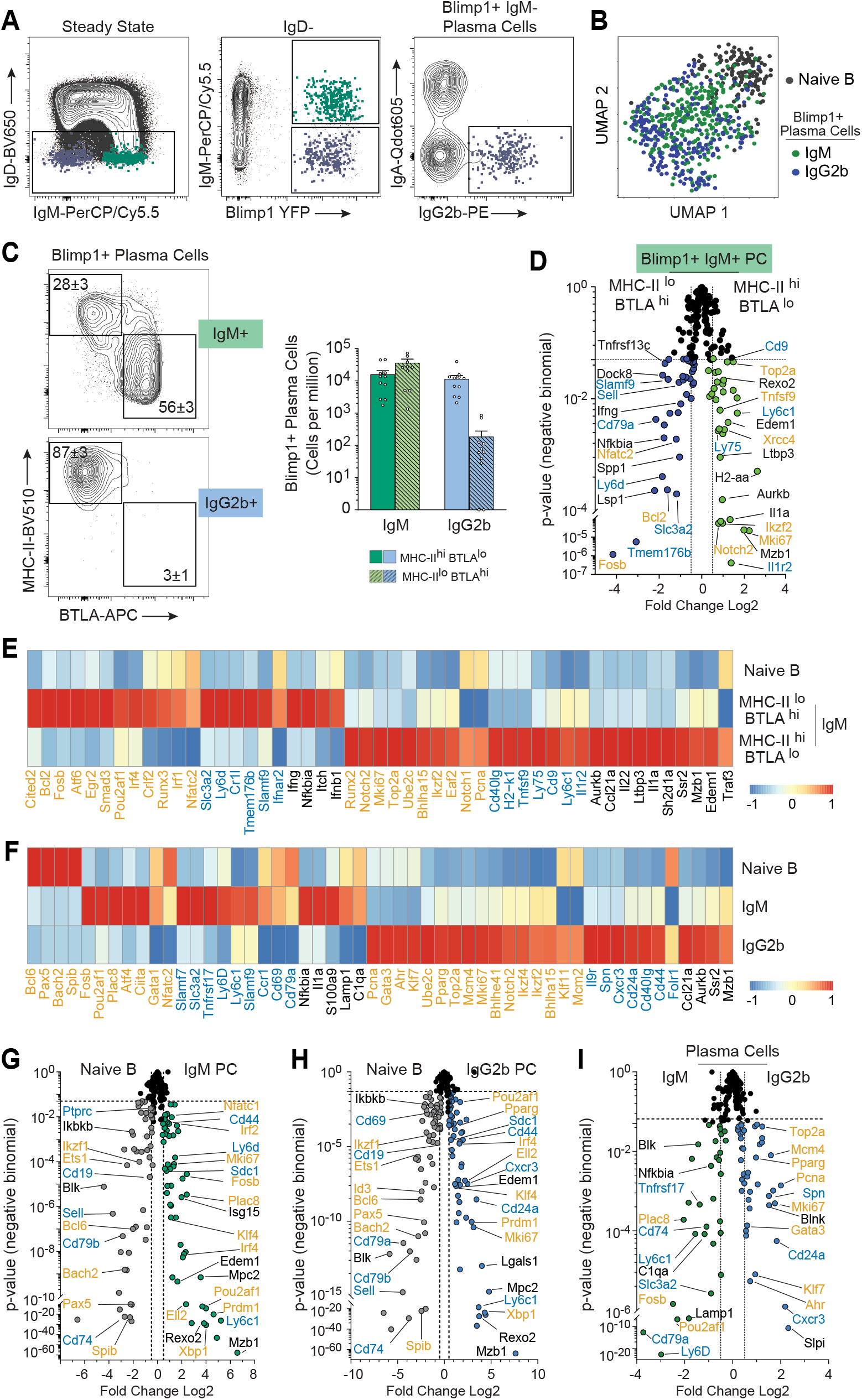
Divergent IgM and IgG2b PC programs in the steady state. (**A**) FACS index sorted IgM^+^ and IgG^+^ PCs taken from SP of an unimmunized Blimp-YFP mouse. (**B**) Dimensional reduction on index sorted PCs from (**A**) based on gene expression distribution using uniform manifold approximation and projection (UMAP). n=303 IgM^+^, 220 IgG2b^+^, 150 naive B cells. (**C**) Expression of surface MHC-II and BTLA by IgM^+^ and IgG2b^+^ PC compartments. Numbers indicate positive percentage expressing. At right, quantification in cells per million of each PC isotype by phenotypic subpopulations (mean ± s.e.m., n=11). (**D**) Differences in gene expression plotted by statistical significance and presented as a volcano plot comparing MHC-II^hi^BTLA^lo^ (n=141) and MHC-II^lo^BTLA^hi^ (n=92) IgM^+^ PCs. (**E**) Pseudobulk heatmap representation of differentially expressed genes generated in **D** compared to naïve B cells, or (**F**) differentially expressed genes between naïve B cells and class specific PCs as shown in volcano plot representations of differentially expressed genes between naïve B cells and (**G**) IgM^+^, (**H**) IgG2b^+^ or (**I**) IgM^+^ versus IgG2b^+^ PCs. For pseudobulk heatmaps, the color scale is based on z-score distribution. Genes labeled in orange represent transcriptional modifiers translocating to or residing in the nucleus, genes labeled in teal represent surface/membrane genes and genes in black represent cytosolic and secreted genes.

In contrast to MHCII-expressing TD antigen-specific IgG^+^ and IgM^+^ PC (**Fig 3F**), steady-state IgM^+^ PCs presented across two phenotypically distinct compartments with the majority expressing high surface levels of BTLA (B and T lymphocyte attenuator) with low to negligible MHC-II (**Fig 6C; Supplementary Fig 5C**). The BTLA^hi^ IgM subset upregulated a divergent set of transcriptional regulators (*e.g., Irf1, Irf4, Runx3, Crlf2, Bcl2, Fosb, Atf6, Egr2, Smad3*, *Nfatc2*) that contrasted significantly to the MHC-II^hi^BTLA^lo^ subset (e.g., *Ikzf2, Runx2, Notch 1, Notch2, Bhlha15, Mki67, Pcna, Top2a, Ube2c, Eaf2*) (**Fig 6D-E**). These transcriptional differences extended to cell surface molecules (e.g., *Ly6d, Slamf9, Slc3a2, Tmem176b, Ifnar2* in BTLA^hi^*; Ly6c1, Ly75, Tnfsf9, Cd9* in BTLA^lo^) as well as intracellular and secreted cellular components (*Itch, Ifnb1*, *Ifng* in BTLA^hi^; *Mzb1, Edem1, Aurkb, Il22* and *Il1a* in BTLA^lo^). These differences demonstrate separable control over a wide spectrum of potential IgM^+^ PC effector functions for these two subsets with further transcriptional divergence from recently formed antigen-specific effector IgM^+^ (**Supplementary Fig 5D**).

IgG2b^+^ PC from the steady state shared many transcriptional features seen in the NP-specific IgG2b^+^ response (**Supplementary Fig 5E**). However, significant IgG2b^+^ PC transcriptional differences were seen across all classes of cell function regulators (nuclear, cell membrane, intracellular and secretory) when contrasted to naive B cells and the broad steady-state IgM^+^ PC programs (**Fig 6 F-I**).

### Distinct IgA PC programming for mucosal immunity

As expected, due to their origins and mucosal targeting function, IgA-expressing PC are most abundant in the Peyer’s patch but can also be found in the BM and the SP at steady-state (**Supplementary Fig 6**). All Blimp-1^+^ IgA PC expressed high levels of both intracellular and surface IgA (**Supplementary Fig 7A**). Using the Blimp1-reporter model, we isolated IgM^neg^ class-switched IgA^+^ PC for gene expression studies (**Fig 7A**) which broadly separated in UMAP clusters based on the full range of gene expression from qtSEQ analysis (**Fig 7B**). In contrast to IgM^+^ PCs, the majority of the IgA^+^ PC cohort expressed intermediate to high levels of MHC-II and lower levels of BTLA (**Supplementary Fig 5C; Fig 7C**). However, IgA^+^ PC maintain high levels of the BCR co-receptor CD79b, akin to IgM^+^ PC and opposite to IgG2b^+^ PC. (**Fig 7D**). IgA^+^ expression of CD98 and PD-L1 was similar to IgM^+^ PCs but had the lowest expression of the BCR co-receptor CD19. Only IgA^+^ PCs expressed a substantial fraction of Ccr9, contrasted by Cxcr3 expression by IgM^+^ and IgG^+^ PCs (**Fig 7D; Supplementary Fig 7B-D**). As all of these protein expression trends extended into the BM derived population of IgA^+^ PCs, this indicated class conserved rather than location driven programming (**Supplementary Fig 7E**). Hence, even at the broad level of surface phenotype, it is clear that IgA^+^ PC express a different molecular program to PC of other antibody isotypes.

**Figure 7:**
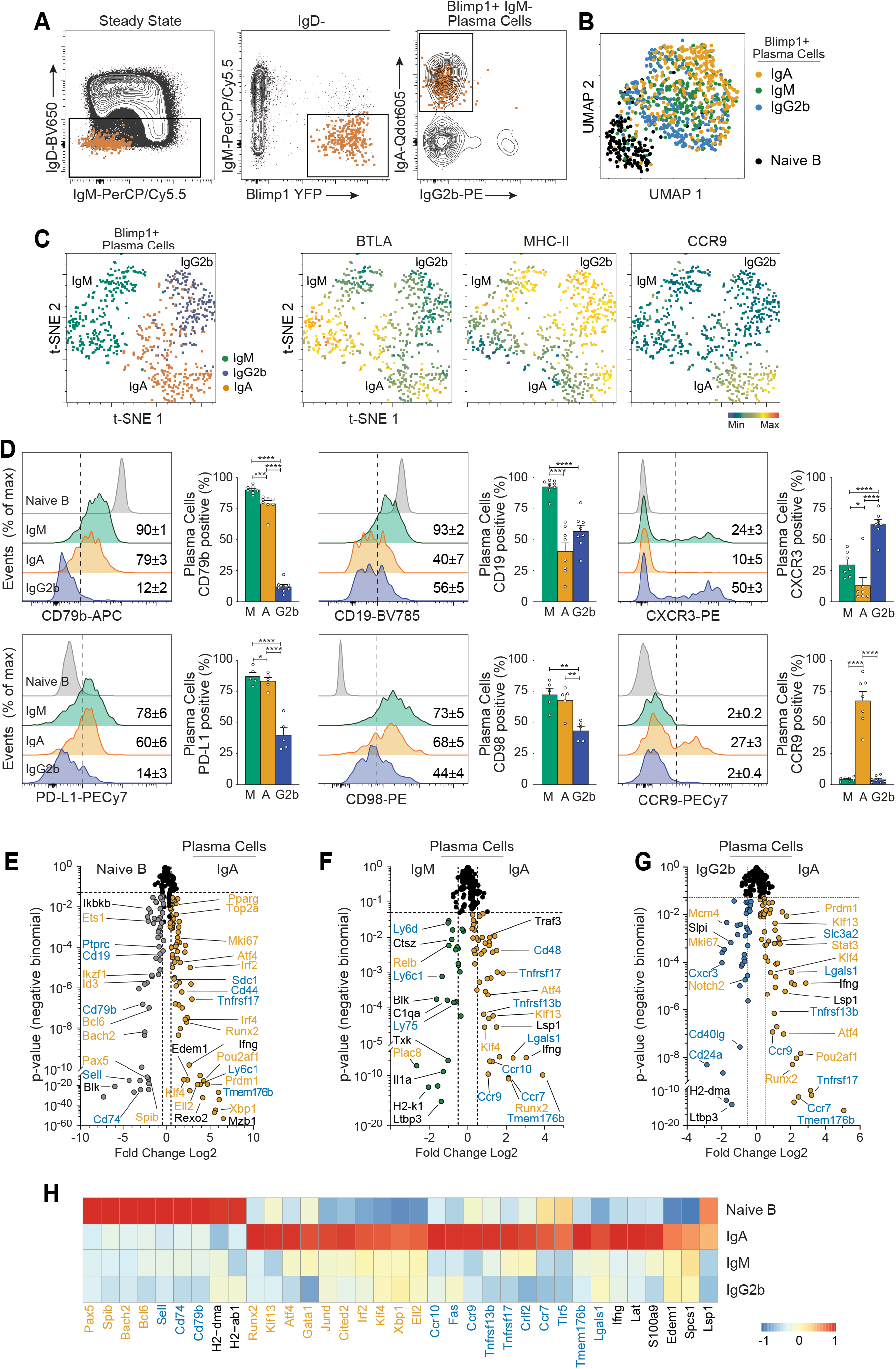
Distinct IgA PC programming for mucosal immunity. (**A**) FACS index sorted splenic-derived IgA^+^ PCs (n=276) taken from spleen (SP) of an unimmunized Blimp-YFP mouse. (**B**) tSNE (t-distributed stochastic neighbor embedding) dimensional reduction on index sorted IgM^+^, IgA^+^ and IgG2b^+^ PCs based on phenotype with overlay of expression level for selected proteins. (**C-D**) Expression of surface protein markers from Blimp-YFP^+^ class specific PCs compared to naïve B cells. Percentage of cells expressing is indicated on plots with quantification at right of percentage expressing (mean ± s.e.m., n = 6; 2 tailed unpaired Students *t-*test, *p < 0.05, ** p<0.01, *** p<0.001, **** p<0.0001). (**E**) Differences in gene expression plotted by statistical significance and presented as volcano plots comparing naïve B cells and IgA^+^ PC, (**F**) IgM^+^ versus IgA^+^ PC and (**G**) IgG2b^+^ versus IgA^+^ PC. (**H**) Pseudobulk heatmap representation of differentially expressed genes focused on IgA^+^ PCs. The color scale is based on z-score distribution. In **E-G**, genes labeled in orange represent transcriptional modifiers translocating to or residing in the nucleus, genes labeled in teal represent surface/membrane genes and genes in black represent cytosolic and secreted genes. For **A, B, E-G**: n=303 IgM^+^, 220 IgG2b^+^, 276 IgA^+^ PCs, 150 Naïve B.

Comparing the IgA^+^ transcriptional program to naive B cells highlighted many of the canonical features of the PC program seen across all PC isotypes (decreased B cell identity: *Pax5, Bach2, Spib, Bcl6*; decreased antigen processing and presentation: *CD74, H2-dma, H2-ab1*; increased lineage transcription factors: *Prdm1, Irf4, Pou2af1*; increased *Sdc1, Ly6c1, CD44, Mzb1, Rexo2*) (**Fig 7E; Supplementary Fig 7F**). However, multiple molecular features were exaggerated in the steady-state IgA^+^ PC compartment in contrast to IgM^+^ and IgG2b^+^ such as the increased expression of major transcriptional regulators (*Runx2, Klf13, Gata1, Jund*), TGF-β responsive transcription factors (*Klf4, Cited2*), UPR and secretion modifiers (*Atf4*, *Xbp1, Ell2, Irf2*) (**Fig 7 F-G**). That IgA^+^ PCs exhibited differential expression of chemokine receptors (*Ccr7, Ccr9, Ccr10*), the highest levels of growth factor receptors (*Tnfrsf13b, Tnfrsf17*), surface receptor sensors (*Tmem176b, Lgals, Tlr5*) and secretory environmental modifiers (*Ifng, S100a9*) indicating the capacity for enhanced chemotaxis, survival, and environmental sensing in the mucosal microenvironment over IgM^+^ and IgG2b^+^ PCs. Expression of these divergent phenotypic and transcriptional programs may uniquely and differentially impact IgA^+^ PC localization, survival, and effector functionality.

## DISCUSSION

In these studies, a targeted single cell RNA sequencing platform connected surface phenotype with unique transcriptional programming of class specific PCs. Divergent transcriptional programming defined early forming IgM^+^ from IgG^+^ subclass PCs which indicated different effector roles. By comparing effector and post-GC PC cohorts, we highlighted the acquisition of unique molecular programs resulting from T_FH_ cell directed CSR and GC cycling necessary for memory phase survival and function. Extension into steady state immunity supported the broad IgM^+^ and IgG^+^ PC transcriptional divergence seen in effector PCs yet revealed two transcriptionally distinct subpopulations within IgM^+^ PCs. Interrogation of IgA^+^ PCs revealed programmatic capacity for mucosal protective functions. Taken together, these separable immune response modules for PC transcriptional control emphasize unique class specific molecular programming that shape early effector responses and impact long-live ‘memory’ PC compartments.

Effector PCs are formed rapidly upon antigen exposure and are important for differentially impacting immune responses through class specific antibody binding of Fc receptors(*10–12*). Effector IgM^+^ PCs upregulated genes coding for Ltbp3 and Tgfb2 that form a secreted complex which can induce T_H_17 or regulatory T cell (T_REG_) formation in a concentration dependent manner(*48*), suggesting effector IgM^+^ PCs may be capable of influencing a pro or anti-inflammatory environment. Additionally, IgM^+^ upregulation of secreted costimulatory cytokine IL-1a may impact proinflammatory Il-1R driven signaling. In line with their antibody driven inflammatory function, IgG2a^+^ and IgG2b^+^ PCs each upregulated additional proinflammatory molecules; however, both subclasses also upregulated anti-inflammatory *Slpi* known to prevent runaway proinflammatory activity(*49*). Therefore, we demonstrate effector PCs are transcriptionally poised for separable class linked functionality via additional secreted molecules.

Gene upregulation of inhibitory cell surface proteins (*Ctla4, Slamf7*) by IgM^+^ PCs would permit T cell modulation during T cell cognate contact, while IgG2a^+^ PC elevated expression of *Tnfsf9* (CD137L), involved in bidirectional signaling after cognate contact, may direct increased inflammatory activity(*50*). As all effector PCs retained MHC-II expression, they may participate in T cell cognate interactions and induce additional T_H_ cell functions through MHC-II antigen presentation as we have previously demonstrated in class switched post-GC PCs(*38*). Thus, each class of effector PC may be capable of impacting ongoing immune responses through differentially impacting and influencing T_FH_ cell functionality.

The GC reaction is critical for generating antigen specific immunological memory through the formation of memory B cells and post-GC PCs poised for long term functionality in specialized survival niches. During GC cycling, higher affinity B cells receive increased T cell help(*51, 52*) and are more likely to exit as PCs(*31–33, 53–55*). Additional T cell interactions may be responsible for imprinting unique transcriptional programs in post-GC PCs. After transit to the BM, PCs exhibit transcriptional and phenotypic changes(*29, 56–58*). However, upon GC exit it is already apparent that post-GC PCs are broadly and indelibly changed to accommodate their ‘memory’ role. For example, they lose cell cycle and division programs that allow entry into a quiescent state and negative regulators of apoptosis for long term survival. Post-GC PCs upregulated of extrinsic signal receptors, costimulatory molecules, and cell adhesion regulators. The effector molecules could contribute to enhanced T cell interaction during GC cycling and may be important for survival niche maintenance via exogenous secreted survival signals and extrinsic contact from support cells such as T_REGs_(*59*).

The steady state IgM^+^ PC compartment phenotypically split into two subsets defined by BTLA^hi^MHC-II^lo^ and BTLA^lo^MHC-II^hi^ expression. As mentioned earlier, subsets of CD138^+^ IgM^+^ PCs have been shown to secrete IL-10 and IL-35 which suppress effector CD4^+^ cells and innate cells to enact a regulatory B cell (B_REG_) role(*19, 60, 61*). The BTLA^hi^MHC-II^lo^ IgM^+^ compartment identified here exhibited shared programs with regulatory IgM^+^ PCs, such as higher expression of *Irf4*(*19*), downregulation of cell cycle linked genes, reduced MHC-II protein and expression of BLTA (associated with *Btla* gene upregulation in regulatory IgM^+^ PCs)(*60*). In contrast, the high protein expression of MHC-II by the second IgM^+^ subpopulation was more similar to helper T cell driven antigen-specific PCs and steady state IgG^+^ PCs. This BTLA^lo^MHC-II^hi^ IgM^+^ compartment upregulated antigen processing and presentation programs, T cell costimulatory proteins (e.g., *Cd40lg, Tnsf9*) and signal molecules, which may indicate capacity for modulating T_H_ cell function during MHC-II antigen presentation as described previously(*38*). Thus, we propose that the phenotypically defined subsets represent transcriptionally distinct IgM^+^ PC cohorts: a B regulatory compartment and a T cell interacting compartment primed for antigen presentation that is TD in origin.

In the steady state, T cell dependent gut derived antigens generate IgA^+^ B cells that can enter GCs to undergo affinity maturation and exit as post-GC PCs(*39, 62–65*). Expression of PD-L1 by IgA^+^ PCs may be retained from GC cycling where PD-1/PD-L1 interactions are critical for cell survival and post-GC PC formation(*66*). Additionally, PD-L1^+^MHCII^+^ IgA^+^ PCs in the lamina propria induced FoxP3^+^ T_REG_ formation in the presence of TGF-β(*67*) which are important for maintaining T_H_17 and IgA^+^ PC homeostasis(*68*). We detected IgA^+^ PC upregulation of many complementary molecular programs with T_H_17 cells, such as transcription factors (*Klf4, Cited2*), and surface protein expression (e.g. *Tmem176b*(*69*), *Fas*(*70*), and chemokine receptors *Ccr7, Ccr9, Ccr10*(*71*)) that may confer similar mucosal defense functionality. Though T_H_17 cells can induce formation of IgA^+^ PCs(*72*), upregulation of *Ifng* by IgA^+^ PCs may in turn allow modulation of T_H_17 formation and function through secretion of IFN-γ(*73*). These reciprocal molecular programs of T_H_17 and IgA^+^ PCs may result from shared formation in a TGF-β rich environment prior to IgA^+^ PC survival niche localization in the spleen(*74, 75*), similar to previous reports that gut IgA^+^ PC translocate to bone marrow survival niches for systemic antibody secretion(*76–78*).

Previous studies on humans have identified preferential and sequential switching capacity of B cells(*79*) and identified class linked heterogeneity which may influence B cell fate and function(*80–82*). As each murine antibody class investigated in this study has a homologous isotype in humans, this work is applicable to ongoing human research, particularly for PC driven diseases, immunotherapeutics and vaccine design. Programmatic studies in Multiple Myeloma have led to approval of the anti-Slamf7 drug Elotuzumab(*83*) and proposals for Hif1a suppression as therapy(*84*); our studies identified upregulation of *Slamf7* in IgM^+^ PCs and *Hif1a* in IgG2b^+^ PCs. Therefore, further exploration of the class specific molecular programs in PCs can potentially identify targetable surface molecules or transcriptional pathways pertinent to PC directed diseases.

Chimeric antigen receptor (CAR) T cells marked the advent of synthetically engineered immunotherapeutics, which has now expanded into B cells(*85–87*). The specificities of known neutralizing antibodies introduced to BCR gene loci generated target-specific antibodies without prior antigen encounter, GC cycling or affinity maturation after PC induction(*85–87*). Beyond specificity, these ‘synthetic’ PCs need to function and survive long term in appropriate niches. Our studies detailing the molecular programming of post-GC ‘memory phase’ PCs provides elements of an isotype-specific transcriptional roadmap for future studies. Previous work has indicated B cell class may impact GC entry(*81*), or terminal differentiation pathway into memory B or PC(*88*). Here our study emphasizes that B cell class may impact immune function beyond Fc binding. We propose that each class of PC is transcriptionally primed to differentially participate in and impact ongoing immune responses via ancillary effector functionality that must be considered during vaccine design.

## MATERIALS and METHODS

### Mice

C57BL/6 (B6), B6.CD45.1 (B6.SJL-Ptprc^a^Pepc^b^/BoyJ) and Blimp1-YFP mice (provided by Susan Kaech, Salk Institute) were bred and housed in specific pathogen-free conditions. All experiments were done in compliance with federal laws and institutional guidelines as approved by The Scripps Research Institutional Animal Care and Use Committee.

### NP-KLH Immunization

Mice were given primary immunization subcutaneously at the base of the tail with 400µg NP-KLH (4-hydroxy-3-nitrophenylacetyl (Biosearch) conjugated to keyhole limpet hemocyanin (Pierce)) mixed with adjuvant based on Monophosphoryl Lipid A.

### Flow Cytometry

Single cell suspensions of draining (inguinal and periaortic) LN, mesenteric LN, Peyers patches (combined), SP and bone marrow (from 2 femurs and 2 tibias) were prepared, followed by incubation in ACK lysis buffer (Gibco) for lysis of red blood cells. Cells were resuspended in phenol red-free DMEM (Gibco) with 2% (vol/vol) FBS at a density of 4 × 10^8^ per ml for staining. Antibody to CD16/32 (2.4G2; produced ‘in-house’) was first added for 10 min on ice, before the addition of ‘cocktails’ of fluorophore-labeled or biotin-labeled monoclonal antibodies (monoclonal antibodies used in **Supplemental Table 1**) prepared in brilliant stain buffer (BD) ‘cocktail’ at pre-titrated quantities, followed by incubation for 30 min on ice. After samples were washed, biotin-labeled antibodies were detected by incubation for an additional 30 min.

For surface staining of IgM, IgG1, IgG2b, or IgA, non-specific binding was blocked with anti-CD16/32 (identified above) for 10 minutes on ice prior to incubation for 30 min on ice with anti-mouse Ig antibodies (**Supplemental Table 1**). Cells were washed and nonspecific binding was blocked by incubation with 2% (vol/vol) normal mouse and normal rat serum followed by incubation with pre-titrated immunoglobulin specific antibody cocktails prepared in brilliant stain buffer (BD) for 30 min on ice. Cells were washed and resuspended in a solution of phenol red-free DMEM (Gibco) with 2% (vol/vol) FBS.

For intracellular staining of Blimp-1, Ki-67, and internal IgA, the surfaces of cells were stained as described above, and cells were labeled with eFluor 780 viability dye (eBioscience). Then cells were fixed, and permeabilized according to the protocol provided in a transcription factor staining kit (eBioscience). Nonspecific intracellular binding was blocked with 2% mouse and rat serum in permeabilization buffer for 5 minutes on ice. Cells were stained for 30 min on ice with pre-titrated immunoglobulin specific antibody cocktails. Cells were washed and resuspended in FACs wash (described above).

Single cells were sorted using a four-laser FACS Aria III with FACS Diva 8.0 software equipped with index sorting software (BD Biosciences) for recording of the “index” phenotypic data for each single cell sorted in each well of a 384-well plate. Flow cytometry data were analyzed with FlowJo software (Version 9.9 and 10.6.2, TreeStar).

### Elispot

96 well PVDF-bottomed plates (Millipore) were washed with filtered PBS prior to coating with NP^23^-BSA (conjugated in-house, 25μg/mL) or goat anti-mouse IgM/IgG/IgA antibody heavy and light chain (25μg/mL); IgM (AF6-78; Biolegend), IgG1 (RMG1-1; Biolegend), IgG2b (RMG2b-1; Biolegend), IgG2a (R19-15; BD bioscience), IgG3 (RMG3-1; Biolegend), IgA (RMA-1; Biolegend). Wells were blocked by incubating DMEM media (Gibco) containing 5% FBS for 1 hour at 37°C. For bulk Elispot experiments, unstained cells suspended at 4×10^6^ cells/mL in DMEM (Gibco) with FBS (5% vol/vol) were added to wells at serially titrated quantities (10×10^4^ to 1.25×10^4^ cells). For single cell sorted Elispot experiments, cells were selectively sorted into 96 well plates using a FACS Aria III with FACS Diva software (BD Biosciences). Plates were incubated at 37°C for a minimum of 14 hours, then wells were then washed prior to addition of horse radish peroxidase (HRP) conjugated goat anti-mouse Fc-specific antibodies for 4 hours at room temperature [1:500 dilution for IgM and IgG subclasses, 1:300 for IgA]. HRP conjugated antibodies were all from SouthernBioTech: IgM, IgG, IgG1, IgG2b, IgG2a, IgG3, IgA. Wells were washed and spots detected by adding a filtered solution of 0.2mg/mL AEC (3-amino-9-ethylcarbazole; Sigma) in 0.1M sodium acetate buffer [pH 5.0] and 0.05% hydrogen peroxide (Sigma). Wells were washed and air dried for a minimum of 24 hours prior to counting the spots by visual inspection under a stereomicroscope.

### qtSEQ single cell RNA sequencing

The single cell sequencing protocol “qtSEQ” has been previously described (*40, 41*) but will be detailed here in brief. Single cells were index sorted into 384 well plates containing 1uL of reverse transcription (RT) master mix using a FACS Aria III with FACS Diva software (BD Biosciences) to maintain protein expression with plate well location. The RT master mix per well consisted of: 0.2μL SuperScript II 5x buffer (Invitrogen), 0.03μL SuperScript II taq (Invitrogen), 0.03 DTT 0.1M (Invitrogen), 0.03μL RNaseOUT (Invitrogen), 0.012μL of 25μM dNTPs (Invitrogen), 0.19μL RNase Free H_2_O and 0.5μL of oligoDT primer [1μM stock] (IDT). Reverse transcription was performed at 42°C for 50 minutes followed by heat inactivation at 80°C for 10 minutes.

For each plate, all 384 wells were pooled following RT, and excess oligoDT primer were removed using ExoSAP-IT express (Affymetrix) following manufacturers protocol prior to buffer exchange and volume reduction using AMpureXP SPRIselect paramagnetic beads (BeckmanCoulter) at a 0.8x bead to 1x library ratio.

A nested PCR approach was used for targeted gene amplification, gene specific primers (available upon request). PCR 1 was performed in a 60μL reaction: 35μL cDNA library in H_2_O, 1.5μL RNaseH (5000U/mL, NEB), 1.2μL Phusion DNA polymerase (2000U/mL, NEB), 12μL 5X Phusion HF Buffer (NEB), 0.85μL of 25μM dNTPS (Invitrogen), 1.2μL of 100μM RA5-overhang primer (custom made, 5’AAGCAGTGGTGAGTTCTACAGTCCGACGATC 3’) and 25nM final concentration for each specific gene targeting external primer using the following thermocycling conditions: 37°C for 20 minutes, 98°C for 3 minutes, followed by 10 cycles of 95°C for 30 seconds, 60°C for 3 minutes, 72 °C for 1 minute and final elongation at 72°C for 5 minutes. Removal of excess primers and volume reduction was performed again using AMpureXP SPRIselect beads, at a 0.9x bead to 1x library ratio.

PCR2 was performed in a 20μL reaction: 10μL PCR1 reaction elute, 0.4μL Phusion DNA polymerase (2000U/mL, NEB), 4μL 5X Phusion Buffer (NEB), 0.3μL of 25μM dNTPs (Invitrogen), 2μL of 20μM RA5 (custom made, 5’ GAGTTCTACAGTCCGACGATC 3’), and 25nM final concentration for each specific gene targeting internal primer using the following thermocycling conditions: 95°C for 3 minutes followed by 10 cycles of 95°C for 30 seconds, 60°C for 3 minutes, 72 °C for 1 minute and final elongation at 72°C for 5 minutes. Removal of excess primers and volume reduction was performed again using AMpureXP SPRIselect beads, at a 0.9x bead to 1x library ratio.

PCR3 was in a 20μL reaction: 10μL PCR2 reaction elute, 0.4μL Phusion DNA polymerase (2000U/mL, NEB), 4μL 5X Phusion Buffer (NEB), 0.3μL of 25μM dNTPs (Invitrogen), 2μL of 10μM RP1 primer (Illumina), 2μL of 10μM RPI library specific primer (Illumina) and 1.3μL water using the following thermocycling conditions: 95°C for 3 minutes followed by 8 cycles of 95°C for 15 seconds, 60°C for 30 seconds, 72 °C for 30 seconds. and final elongation at 72°C for 5 minutes.

Removal of excess primers and volume reduction was performed again using AMpureXP SPRIselect beads, at a 0.7x bead to 1x library ratio and the final library eluted in 20μL H_2_O for quantification and library sequencing preparation.

cDNA library concentration and quality were assessed using an Agilent 2100 Bioanalyzer (Agilent Technologies) and a Qubit™ 4 Fluorometer (Invitrogen Technologies). Separately labeled cDNA libraries were multiplexed according to the manufacturer’s protocol (Illumina, San Diego CA). The libraries were sequenced on an Illumina NextSeq500 sequencer using a paired end sequencing run set as: Read 1: 15 cycles; Index Read 1: 6 cycles; Read 2: 71 cycles.

### Single-cell RNA-sequencing analysis

Bcl2fastq (Illumina) was used to demultiplex sequencing libraries and assort reads by well barcodes prior to alignment using Bowtie2 to a custom genome consisting of the amplicon regions for the specifically targeted genes used in the experiment based on the murine genome GRCm38.p6, version R97 (Ensembl). HTseq was used for read and UMI reduced tabulation. Quality control discarded wells that contained fewer than 125UMI counts and fewer than 25 unique genes and removed genes that were not expressed in at least 10 different cells. A secondary filter included naïve B cells wells containing 2 of 3 genes (*Cd79a, Cd79b, Ptprc*) and PC wells containing 2 of 3 genes (*Prdm1, Irf4, Xbp1*).

Seruat (Version 3(*89*)) was used for downstream analysis where raw UMI counts were scaled and normalized through log transformation. The Seurat function “FindMarkers” (“negbinom” distribution) was used to find isotype specific differentially expressed genes and Prism V9.0 software (GraphPad) was used to generate volcano plots. The Seurat function “RunUMAP” (using variable features for clustering) was used for transcription based UMAP clustering. The Seurat function “VlnPlot” was used for generation of violin plots. Differentially expressed gene lists underwent pathway gene ontology (GO) enrichment analysis using PANTHER (version 14.0(*90*)). Pseudobulk heatmaps were generated using the average gene expression after scaling and normalization for each population using the R function “Pheatmap” (scaling by row enabled).

Integration of surface phenotype data using FlowJo (TreeStar) with transcriptional data from qtSEQ matched well barcodes with index sorting location for UMAP clustering on combined phenotype and transcriptional data using Biovinci (BioTuring Inc., San Diego, CA, USA). The Flowjo function “tSNE” was used for tSNE plot generation using phenotypic data.

### Statistics

The statistical tests and methods, number of samples, and number of cells or genes are indicated in the figure legends. Statistics computed with R software detailed above. Mean values, s.e.m. values, and unpaired Students t-tests were calculated and graphed with Prism v9.0 software (GraphPad). A P value of less than 0.05 was considered statistically significant.

## Code and Data availability

All customized R codes used for data analysis and raw and processed data files for the scRNAseq analysis in the study will be provided upon request without restriction.

## Acknowledgements

This work was supported by the US National Institutes of Health (AI047231, AI040215 and AI071182) and Bill & Melinda Gates Foundation (BMGF 0PP1154835) to M.G.M.-W.

## Author Contributions

B.W.H, A.G.S, L.J.M-W, and M.G.M-W designed and developed single cell qtSEQ, B.W.H, A.G.S, K.B.M, A.M.R, L.J.M-W, and M.G.M-W designed and performed experiments, B.W.H, L.J.M-W, and M.G.M-W designed the study, analyzed data and wrote the paper.

---

**Supplementary Fig 1:**
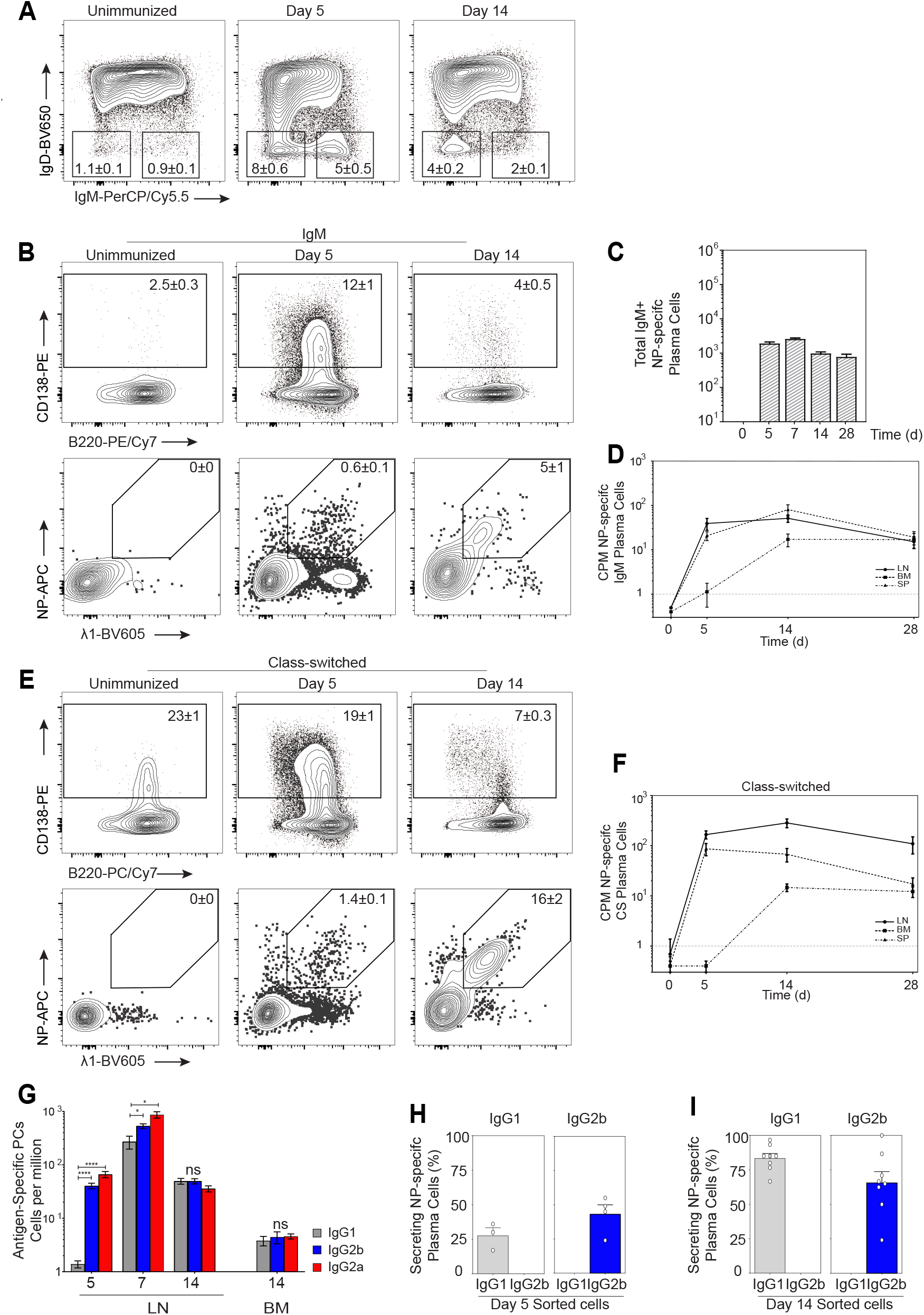
Spatiotemporal access to class specific Plasma Cells. (A) Flow cytometry from draining lymph nodes before (unimmunized), and 5 or 14 days after primary immunization with NP-KLH in MPL adjuvant gated as Gr1-CD3-CD19+ and/or CD138+ B cells. Numbers in outlined areas indicate percent of cells (mean± s.e.m.) (B) Gating strategy of IgM PCs to isolate NP+Lambda1+ PCs with (C) quantification and (D) distribution over time in the dLN (LN), spleen (SP) and bone marrow (BM). (E, F) Gating strategry of Class Switch PCs and temporal quantification of NP+Lambda1+ PCs across dLN, SP and BM at time points indicated after immunization. (G) Quantification of isotype specific NP+Lambda1+ PCs at time points indicated after immunization in the dLN and BM (* p <0.05, **** p<0.0001; 2 tailed unpaired Students t test). Single cell sorted antigen specific Elispot at day 5 (H) and d14 (I) for IgG2b+ (red) and IgG1+ PCs (grey) from the dLN. Frequency is shown after exposure to either anti-IgG2b+ secondary antibody (left column) or anti-IgG1+ secondary antibody (right column). Data are representative of at least two experiments (except day 7 in (C) 1 experiment); mean ± s.e.m. of n = 3-12 mice.

**Supplementary Fig 2:**
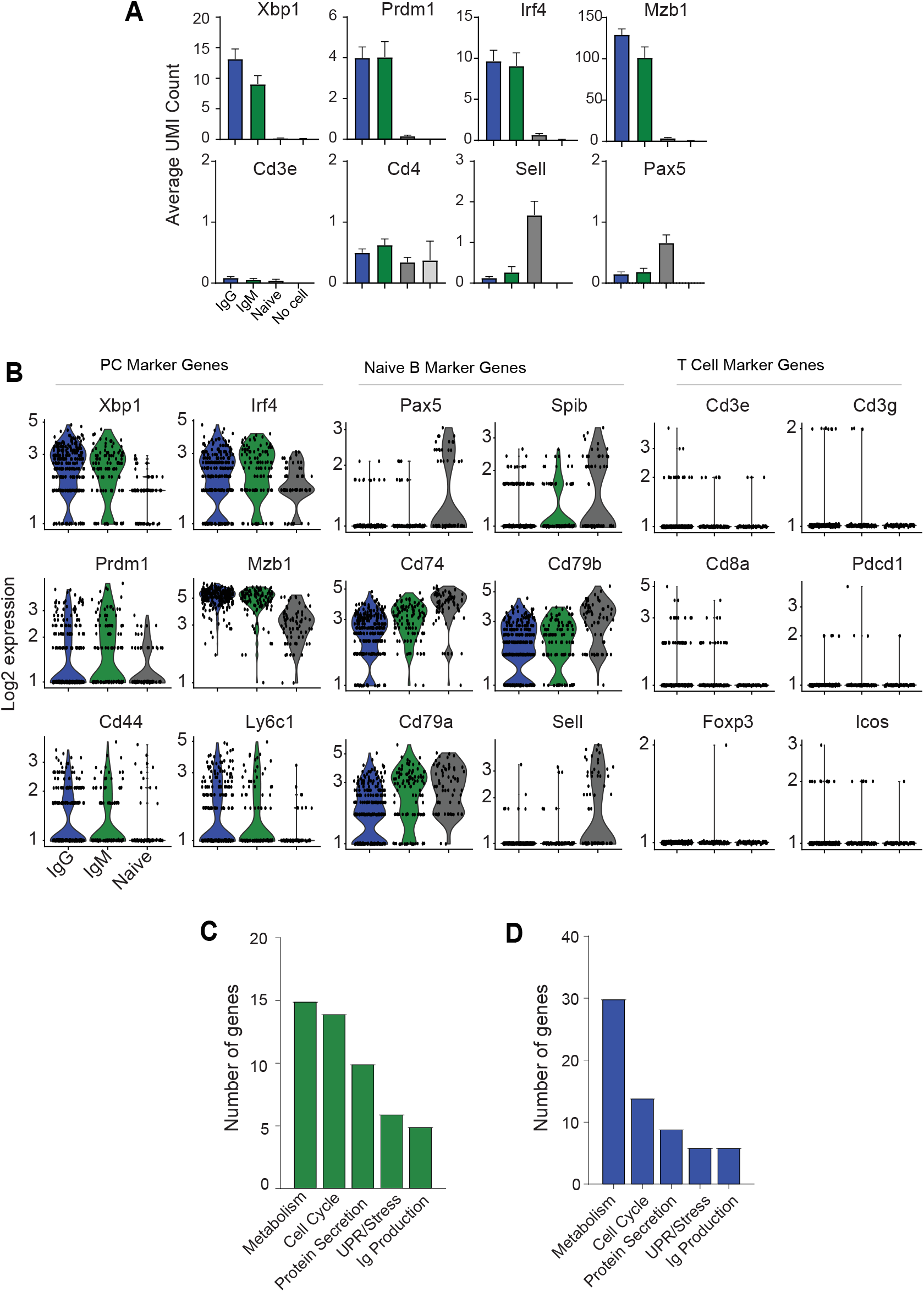
Gene targeted single cell RNA sequencing using qtSEQ. (A) Average expression of raw UMI counts for selected genes taken from IgG, IgM, Naive B single cells compared to control empty well samples. n=109 IgM+, n=269 IgG+, n=73 naive B cells, n=16 no cell control wells. (B) Normalized and scaled UMI counts plotted using a Log2 scale for PC, Naive B and T cell lineage marker genes for IgG+, IgM+ and naive B cells used in Figure 2. (C) Go enrichment of the genes upregulated in IgM+ PCs over naive B cells or (D) IgG+ over naïve B cells. (Protein Metabolism, GO:0051246; Cell Cycle, GO:0051726; Protein Secretion, GO:1903530; UPR/Stress, GO:0006986; Immunoglobulin Production, GO:0002639).

**Supplementary Fig 3:**
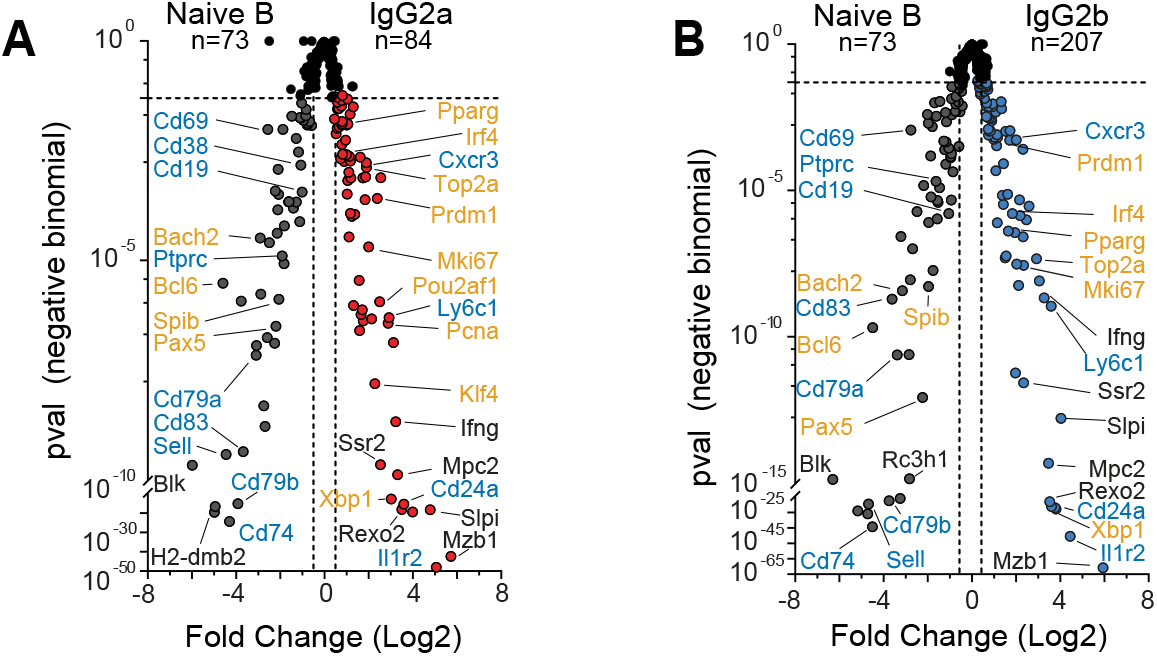
Distinct programming of inflammatory subclass IgG effector PCs compared to naïve B cells. (A) Differences in gene expression plotted by statistical significance and presented as volcano plots comparing naïve B cells and day 5 antigen specific IgG2a+ or (B) IgG2b+ PC. For index sorted cells n=207 IgG2b+, n=84 IgG2a+ PCs, n=73 naïve B cells. Genes labeled in orange represent transcriptional modifiers translocating to or residing in the nucleus, genes labeled in teal represent surface/membrane genes and genes in black represent cytosolic and secreted genes.

**Supplementary Fig 4:**
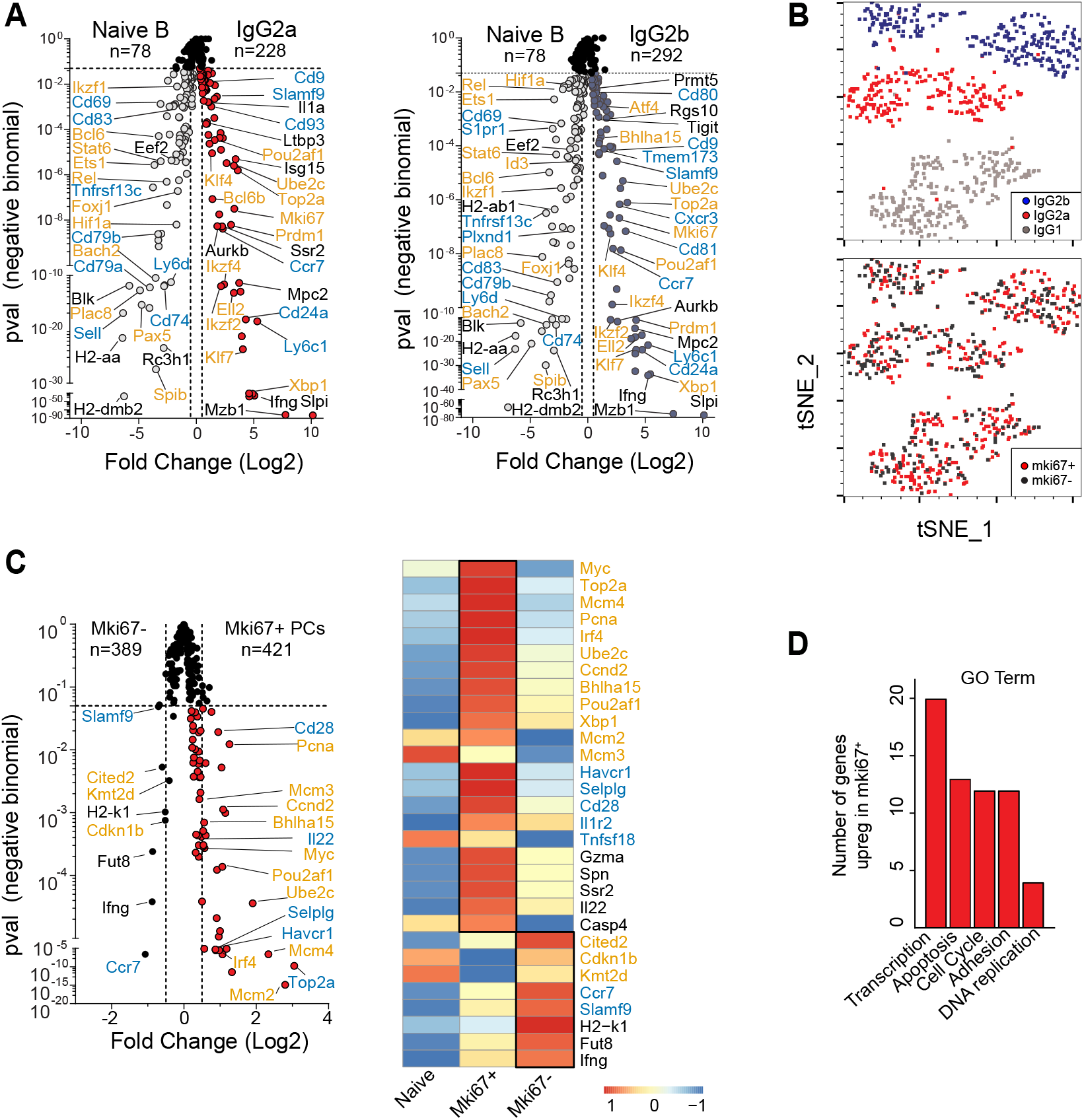
Time from GC cycling impacts post-GC PC transcriptional programming. (A) Differences in gene expression plotted by statistical significance and presented as volcano plots comparing naive B cells (CD19+IgD+IgM+ Lambda1+CD138-NP-Gr1-CD3e-) and antigen specific IgG2a+ or IgG2b+ PCs. (B) Dimensional reduction based on surface phenotype of index sorted PCs using tSNE (t-distributed stochastic neighbor embedding) with positive expression of Mki67 overlaid on cells in red. (n=290 IgG1+, n=292 IgG2b+, n=228 IgG2a+ PCs). (C) Differences in gene expression plotted by statistical significance and presented as a volcano plot comparing Mki67+ and Mki67-PCs with pseudobulk heatmap representation of differentially expressed genes. The color scale is based on z-score distribution. (D) Number of genes for each GO enrichment category of genes upregulated by Mki67+ PCs. (Transcription, GO:0045944; Apoptosis, GO0043065; Cell Cycle, GO:0051726; Adhesion, GO:1903037; DNA replication, GO:0006261).

**Supplementary Fig 5:**
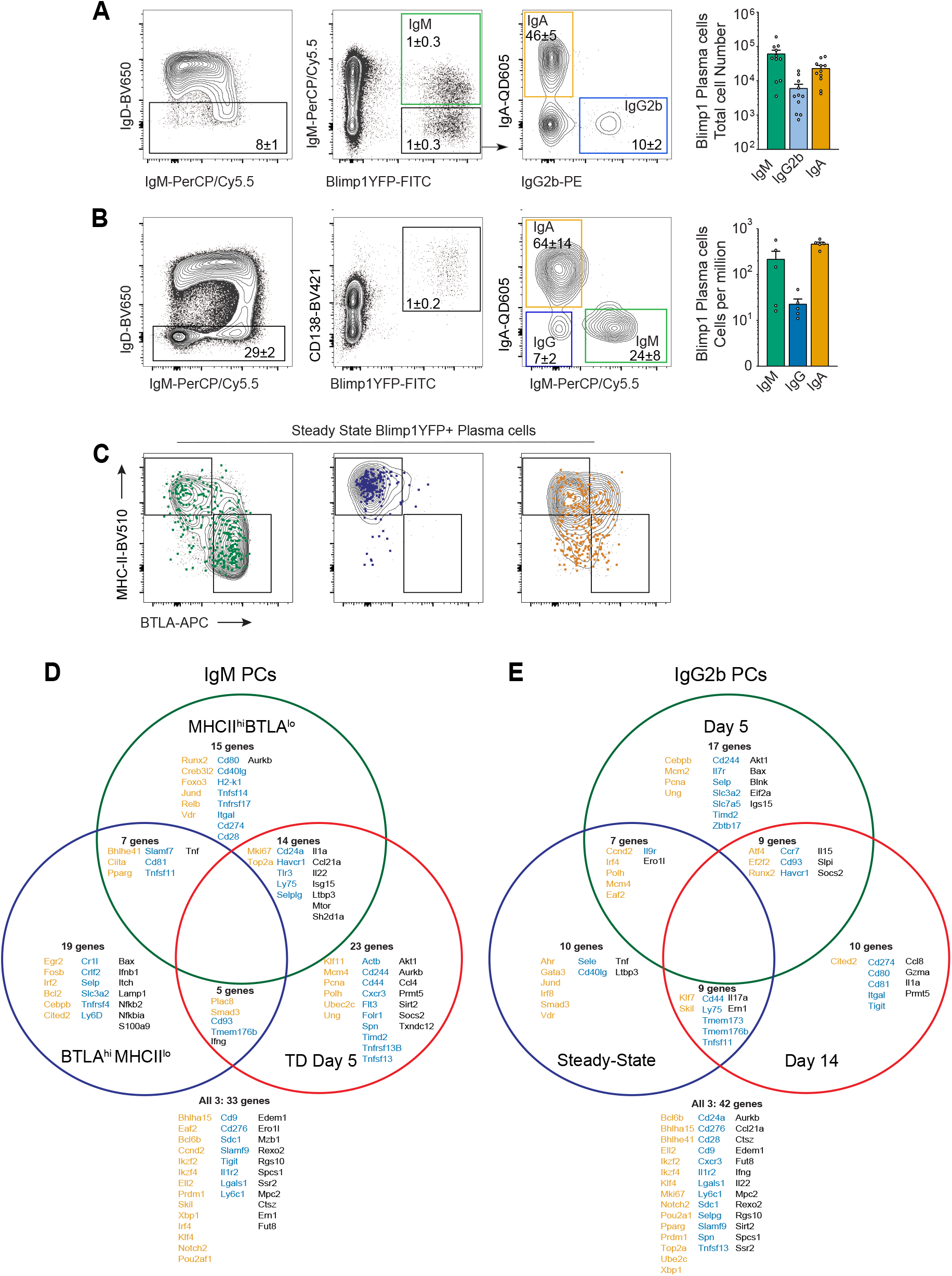
Conservation of class-linked molecular programming of PCs. (A) Representative FACS gating strategy for isolation of IgM+, IgG2b+, and IgA+ PCs taken from spleen (SP) of an unimmunized Blimp-YFP mouse. Quantification of total PCs by class at right (mean ± s.e.m., n=11) (B) Representative FACS gating strategy for isolation of IgM+, IgG+, and IgA+ PCs taken from BM of an unimmunized Blimp-YFP mouse. (mean ± s.e.m., n=5). Quantification of PCs by class in cells per million at right. (C) Index sorted PCs distributed by MHC and BTLA protein expression overlaid on bulk population gating. Venn diagram showing genes that are upregulated in each defined IgM+ PC (D) or IgG2b+ (E) PCs compared to naïve B cells. IgM PCs are divided into day 5 effector (n=109), MHC-IIhiBTLAlo (n=141) and MHC-IIloBTLAhi (n=91). IgG2b+ PCs are divided into day 5 (n=185), day 14 (n=292) and steady state (n=220) cells. Overlapping regions indicated shared upregulation of genes. Genes upregulated in all 3 populations are listed outside the venn diagram below. Genes labeled in orange represent transcriptional modifiers translocating to or residing in the nucleus, genes labeled in teal represent surface/membrane genes and genes in black represent cytosolic and secreted genes.

**Supplementary Fig 6:**
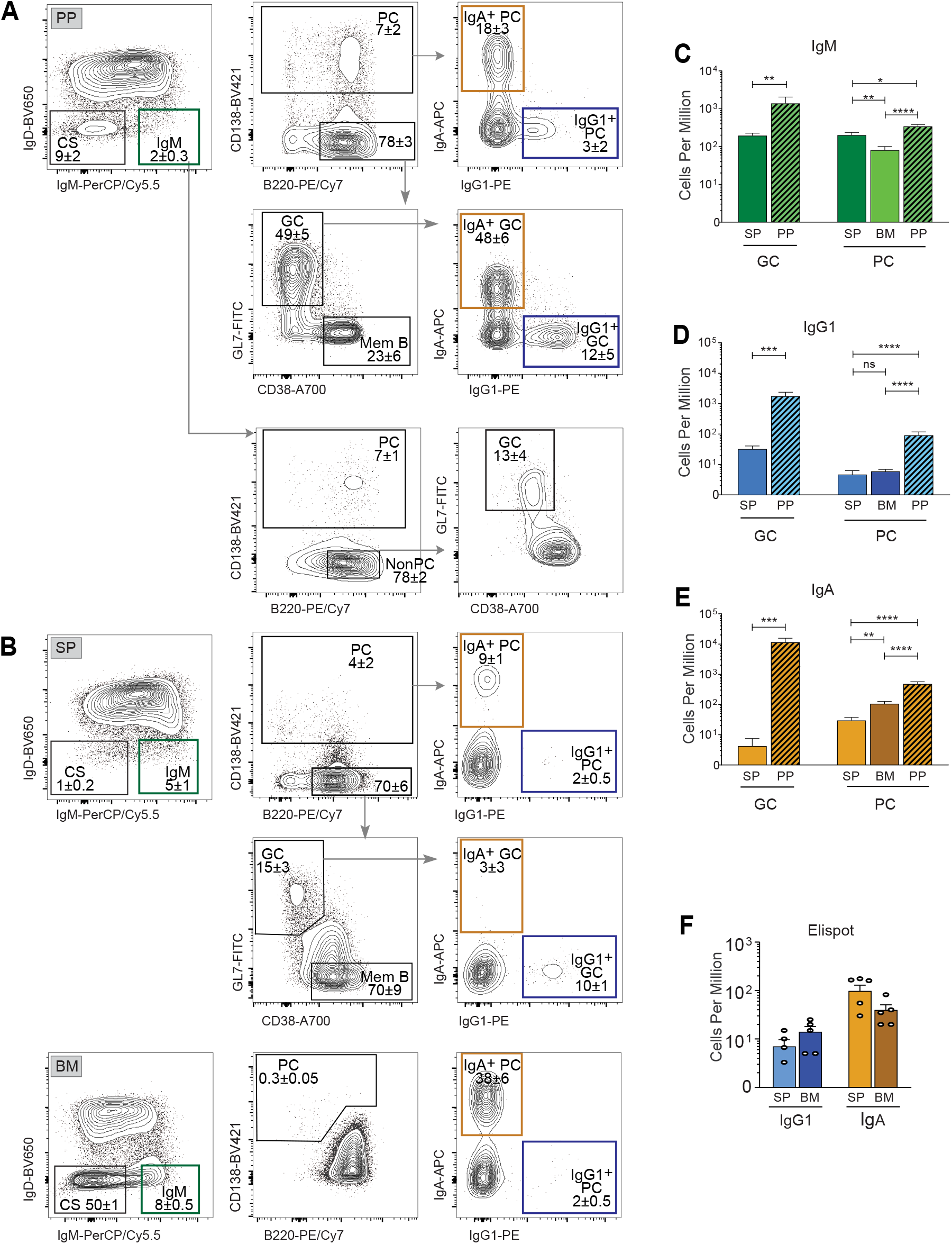
The ongoing response in the steady state murine system presents organ and B cell class based heterogeneity. (A) Representative gating of B cells from the Peyer’s Patches (PP) of an unimmunized C57/B6 mouse divided into class switched (CS, IgD-IgM-) or IgM+ populations. Plasma cells (PC, CD138+) and Germinal center (GC, CD138-GL7+CD38-) cells are shown divided by IgM+, IgA+ and IgG1+ isotypes. (B) Representative gating of B cells from the spleen (SP, top) and bone marrow (BM, bottom) of an unimmunized C57/B6 mouse. Quantification of IgM+ (C), IgG1+ (D), and IgA+ (E) B cells from the SP, PP, BM as gated in a-c. (For PP n=6, SP n=14, BM n=15; mean ± s.e.m., two tailed unpaired t test; *p<0.05, **p<0.01, ***p<0.001, ****p<0.0001). (F) Elispot frequency in cells per million of PCs from the SP and BM (n=5 mean ± s.e.m.).

**Supplementary Fig 7:**
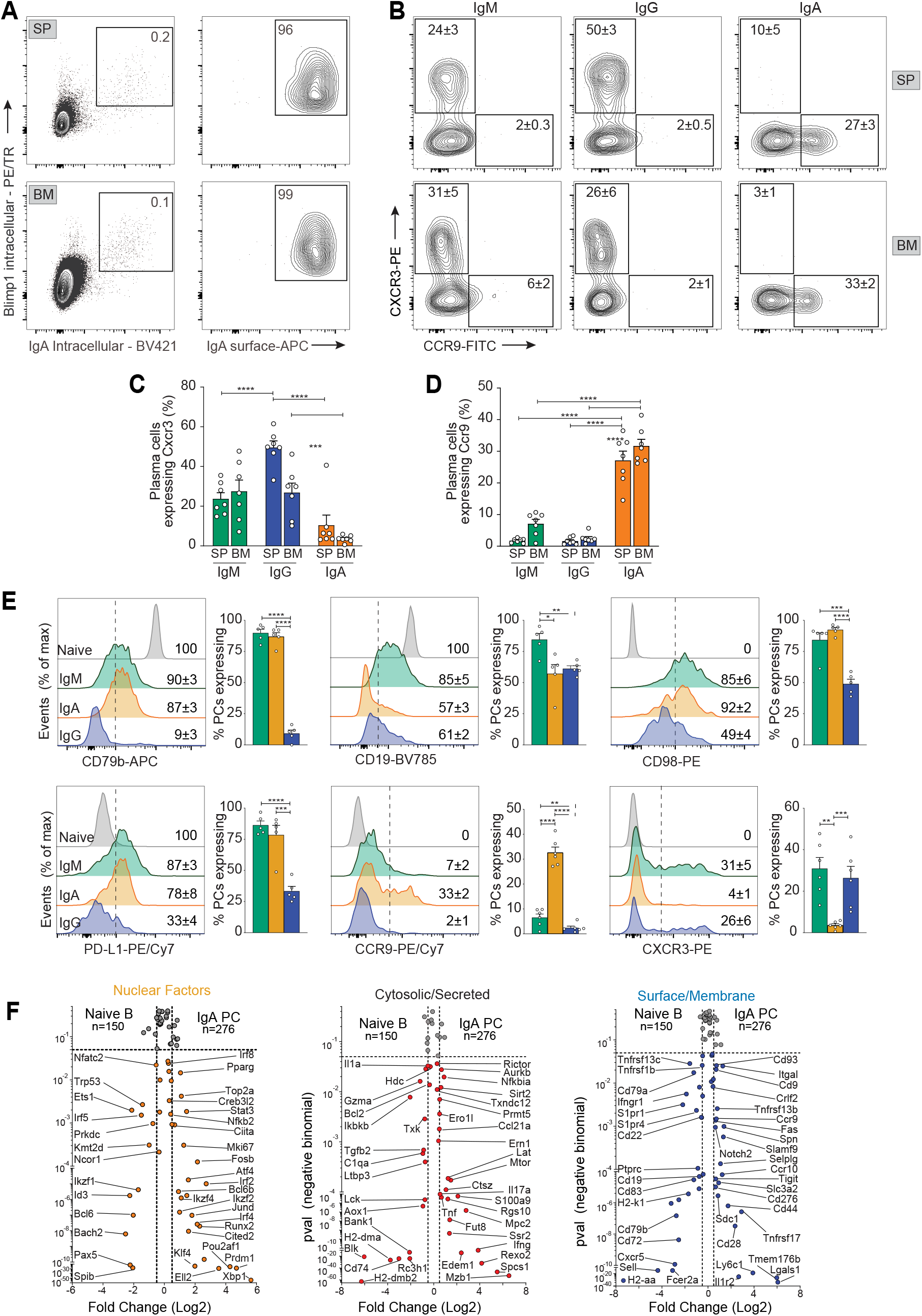
Retention of IgA+ PC programming in the bone marrow. (A) Class switched B cells (Gr1-Cd3e-IgM-IgD-, CD19+ and/or CD138+) expressing both intracellular Blimp-1 and intracellular IgA from the SP (top) and BM (bottom) are shown gated for surface IgA. (B) Representative FACs gating of IgM+, IgG+ and IgA+ PCs from the SP (top) and BM (bottom) of unimmunized C57/B6 mice, divided by surface expression of Cxcr3 and Ccr9. Frequency of PCs of each isotype expressing Cxcr3 is shown in (C) and Ccr9 shown in (D). (For b-d, n=7, mean ± s.em). (E) Expression of surface protein markers from class specific BM PCs compared to naive B cells with percentage of cells expressing listed (mean ± s.e.m. of n=6). All statistics 2-tailed unpaired t test; *p < 0.05, ** p<0.01, *** p<0.001, **** p<0.0001. (F) Differences in gene expression plotted by statistical significance and presented as volcano plots comparing naive B cells (CD19+IgD+IgM+BlimpYFP-CD138-Gr1-CD3e-) and IgA+ PCs separated by gene categorization (n=150 naïve B, n=276 IgA PC).

**Supplementary Table 1:** Antibodies used in flow cytometry.

| **Antibody Target** | **Clone** | **Fluorophore** | **Provider** |
| --- | --- | --- | --- |
| CD79b | HM79-12 | APC | Biolegend |
| NP | - | APC | MMW Lab |
| CD272 (BTLA) | 8F4 | APC | eBioscience |
| IgG1 | X56 | APC | BD Biosciences |
| IgA | RMA-1 | APC | Biolegend |
| Streptavidin-APC | - | APC | Biolegend |
| CD38 | 90 | AlexaFluor700 | eBioscience |
| Efluor Viability | - | eFluor 780 | eBioscience |
| IgG1 | RMG1-1 | APC-Cy7 | Biolegend |
| CD138 | 281-2 | PE | BD Biosciences |
| CD183 (CXCR3) | 173 | PE | Biolegend |
| CD98 | 4F2 | PE | BD Biosciences |
| IgG2b | RMG2b-1 | PE | Biolegend |
| IgG1 | A85-1 | PE | Biolegend |
| Blimp-1 | 5E7 | PE | BD Biosciences |
| B220 | RA3-6B2 | PE-TR | Biolegend |
| Blimp-1 | 6D3 | PE-TR | BD Biosciences |
| GR1 | RB6-8C5 | PE-Cy5 | Biolegend |
| CD3e | 2C11 | PE-Cy5 | Biolegend |
| F4/80 | BM8 | PE-Cy5 | eBioscience |
| PD-L1 | 10f.9g2 | PE-Cy7 | Biolegend |
| B220 | RA3-6B2 | PE-Cy7 | Biolegend |
| CXCR3 | CXCR3-173 | PE-Cy7 | Biolegend |
| CCR9 | CW1.2 | PE-Cy7 | Biolegend |
| IgG2b | R12-3 | FITC | BD Biosciences |
| GL7 | GL7 | FITC | Biolegend |
| Ki67 | 16A8 | FITC | Biolegend |
| Fas | J02 | FITC | BD Biosciences |
| CD79b | HM79-12 | FITC | Biolegend |
| IgM | 331.12 | PerCP-Cy5.5 | MMW Lab |
| CD138 | 281-2 | Brilliant Violet 421 | BD Biosciences |
| IgA | C10-1 | Brilliant Violet 421 | BD Biosciences |
| IgG1 | RMG1-1 | Brilliant Violet 421 | Biolegend |
| Streptavidin- BV421 | - | Brilliant Violet 421 | Biolegend |
| I-A/I-E (MHCII) | M5/114.15.2 | Brilliant Violet 510 | Biolegend |
| Streptavidin-BV510 | NA | Brilliant Violet 510 | Biolegend |
| CD19 | 1D3 | Brilliant Violet 510 | BD Biosciences |
| Streptavidin- QD605 | - | Qdot605 | Invitrogen |
| IgD | 11.26 | Brilliant Violet 605 | Biolegend |
| CD138 | 281-2 | Brilliant Violet 605 | Biolegend |
| IgM | RMM-1 | Brilliant Violet 605 | Biolegend |
| CD19 | 1D3 | Qdot625 | MMW Lab |
| IgD | 11.26 | Brilliant Violet 650 | Biolegend |
| IgM | II/41 | Brilliant Violet 650 | BD Biosciences |
| IgA | C10-1 | Brilliant Violet 711 | BD Biosciences |
| CD19 | 6D5 | Brilliant Violet 800 | Biolegend |
| IgA | RMA-1 | Biotin | Biolegend |
| Lambda-1 | R11-153 | Biotin | BD Biosciences |
| IgG2b | RMG2b-1 | Biotin | Biolegend |

